# NDR1/2 kinases regulate cell polarization and cell motility through Cdc42 GTPase and Pard3 signaling in mammalian cells

**DOI:** 10.1101/2025.10.30.685405

**Authors:** Jun Gu, Jelena Marjanovic, Marjana Tomic-Canic, Fangliang Zhang, Fulvia Verde

## Abstract

Controlling cell polarity and the directionality of cell motility is critical for effective cell migration during wound healing. NDR (nuclear dbf2-related) kinase pathways have roles in cell morphogenesis that are conserved from yeast to humans. Here, we reveal that knockdown of NDR1/2 kinases significantly alters cell size, shape, and the actin cytoskeleton, while reducing migration persistence and impairing cell polarization in wound healing assays. Mechanistically, we find that NDR1/2 kinases regulate the spatial and temporal dynamics of Cdc42 GTPase. Reduced NDR kinase levels increase Cdc42 GTPase activity and disrupt Pard3 subcellular location. NDR kinases phosphorylate Pard3 at Serine144, and overexpressing Pard3 can partially restore wound healing in NDR-depleted cells, an effect lost when Serine144 is mutated. Finally, we determine that NDR1 knockdown significantly impairs wound closure in human skin ex vivo wound healing assays, highlighting NDR kinase physiological importance.

Collectively, this study demonstrates that NDR kinases modulate cell motility and polarization through the control of Pard3 and Cdc42 signaling in human fibroblasts.

## Introduction

Most cells are polarized regardless of their differences in shape or function, and this polarization is essential for embryonic and tissue development (Bera and Loeffler, 2025). The asymmetric organization of cellular components, known as cell polarity, allows cells to sense directional cues and coordinate front-rear migration in response to chemotactic signals (Campanale et al., 2017). Evolutionarily conserved polarity protein complexes play a pivotal role in the regulation of cell polarization, including the partitioning-defective (PAR) complex (Pard3, Par6 and aPKC), which interact with Rho family GTPases such as Cdc42 and Rac1 to modulate cytoskeletal dynamics (Dickinson, 2023; Johansson et al., 2000; Nakayama et al., 2008; Nishimura et al., 2005). Of these, Cdc42, in particular, serves as a central regulator of cell polarity, orchestrating actin polymerization, membrane protrusion, and the establishment of leading and trailing edges during cell migration (Cohen et al., 2018; Patwardhan et al., 2024; Ravichandran et al., 2024). Importantly, persistent cell migration requires synchronized actin treadmilling, formation of leading-edge adhesions, and disassembly at the trailing edge. Therefore, the precise spatial and temporal activation of Rho GTPases is crucial for effective directional migration and wound healing.

Nuclear Dbf2-Related (NDR) kinases, NDR1(AKA STK38) and NDR2(AKA STK38L), are members of AGC kinase family known for their crucial role in cell morphogenesis, polarity, proliferation and migration across species, from yeast to humans (Hergovich et al., 2006; Tamaskovic et al., 2003). For instance, in the fission yeast *Schizosaccharomyces pombe*, NDR kinase such as Orb6 spatially restricts Cdc42 activation to cell tips, ensuring polarized growth (Das and Verde, 2013; Nunez et al., 2016). In mammals, NDR1/2 kinases have emerged as key regulators of centrosome duplication, cell cycle progression, immune response, autophagy, and apoptosis (Cornils et al., 2011; Du et al., 2013; Enomoto et al., 2013; Fukasawa et al., 2015; Hergovich et al., 2009; Hergovich et al., 2007; Joffre et al., 2015; Tang et al., 2015; Vichalkovski et al., 2008; Ye et al., 2020). Recent evidence further implicates these kinases in the control of cell polarity and migration in neuronal and epithelial tissues (Keller et al., 2019; Tang et al., 2015; Yang et al., 2014). Depletion of NDR1/2 decreases cell migration speed in human immortalized bronchial epithelial cells by activating the GEF-H1/RhoB pathway(Keller et al., 2019), and loss of NDR1/2 kinases exhibit defective RhoA activation and polarization in mouse thymocytes cells (Tang et al., 2015). However, their exact functions and mechanisms in fibroblast migration and wound healing are not yet fully known.

The polarity protein Pard3 acts as a scaffold for junctional proteins, which is essential for cell–cell contact assembly and maintenance (Kamakura et al., 2024). Pard3 and Cdc42 interactions with the Par6/aPKC protein complex are mutually exclusive, and atypical protein kinase C (aPKC) shuttles between a Pard3/Par6/aPKC and a Cdc42/Par6/aPKC protein complex (Bera and Loeffler, 2025; Vargas and Prehoda, 2023). Phosphorylation dysfunction causes Pard3 mis-localization, disrupting cell polarity and migration (Yamashita et al., 2020). NDR kinases phosphorylate Pard3 at Ser383 preventing dynein interaction and polarizing Pard3 in mouse neuron axons (Yang et al., 2014). Thus, these observations suggest the idea that NDR kinases may have a role in the control of Cdc42 GTPases function in cell migration.

Fibroblasts are highly dynamic cells that exhibit high proliferative and motile capacity, supporting tissue repair in body’s connective system (Ichim et al., 2018; Talbott et al., 2022). These cells detect signals such as chemotactic cues, migrate to the affected area, remodel the extracellular matrix, and contribute to the restoration of tissue integrity (Cen et al., 2021). In this study, we utilized human foreskin fibroblasts as a model system to investigate the role of NDR kinases in cell movement, identifying Cdc42 and Pard3 as key downstream effectors that regulate cell polarization and migration. Our findings uncover a novel mechanism where NDR kinases control the activity of Cdc42 and the distribution of Pard3 at the cell-cell junctions, modulating the directionality of cell migration. Furthermore, we determined that NDR1 kinase has a crucial role in wound closure using human skin *ex vivo* wound healing assays, highlighting the functional significance of this control in human health.

## Results

### Knockdown of NDR1 and/or NDR2 alters fibroblast morphology by regulating cell size, shape, and actin cytoskeletal organization

To investigate the role of NDR kinases on fibroblasts, human foreskin fibroblasts (HFFs) were stably expressed with shRNA by lentivirus targeting *Ndr1* and/or *Ndr2* (Fig. 1, A-C). Consistent with previous reports in other species (Cornils et al., 2010; Schmitz- Rohmer et al., 2015; Tang et al., 2015), NDR kinases compensate for each other in human fibroblasts. Reduction of NDR1 is compensated by NDR2 in a gene-dose manner (Fig. 1, A-C). NDR kinases were reported to be involved in regulation of cytoskeleton remodeling across species (Bettoun et al., 2016; Emoto et al., 2004; Fang and Adler, 2010; Hergovich, 2013; Joffre et al., 2015; Jonischkies et al., 2024; Klimek et al., 2019; Liu et al., 2017; Madencioglu et al., 2021; Santos et al., 2023; Wu et al., 2013). However, less attention has been paid to test if NDR kinases control the cell cytoskeleton in human fibroblasts. To check whether NDR kinases influence fibroblasts cell morphogenesis, cells grown at low density were fixed with 4% PFA for further examination. An unbiased Opera Phenix Plus High-Content Screening Assay was performed to ascertain whether NDR kinases affect cell cytoskeleton assembly in human fibroblasts (Fig. 1, D-H). Immunofluorescence staining of F-actin by Phalloidin-Alexa 488 suggests that NDR kinases influence cell morphology by altering cell spreading area, shape and actomyosin-cytoskeleton (Fig. 1 D). Cell spreading area was measured on adherent cultures, whereas detached cell mapped area was quantified to evaluate intrinsic cell size without substrate influence. The size of trypsinized, detached cells was assessed and found to be consistent with the spreading size, again suggesting that loss of NDR kinases leads to a decrease in cell size (Fig. 1, I-J).

**Figure 1.**
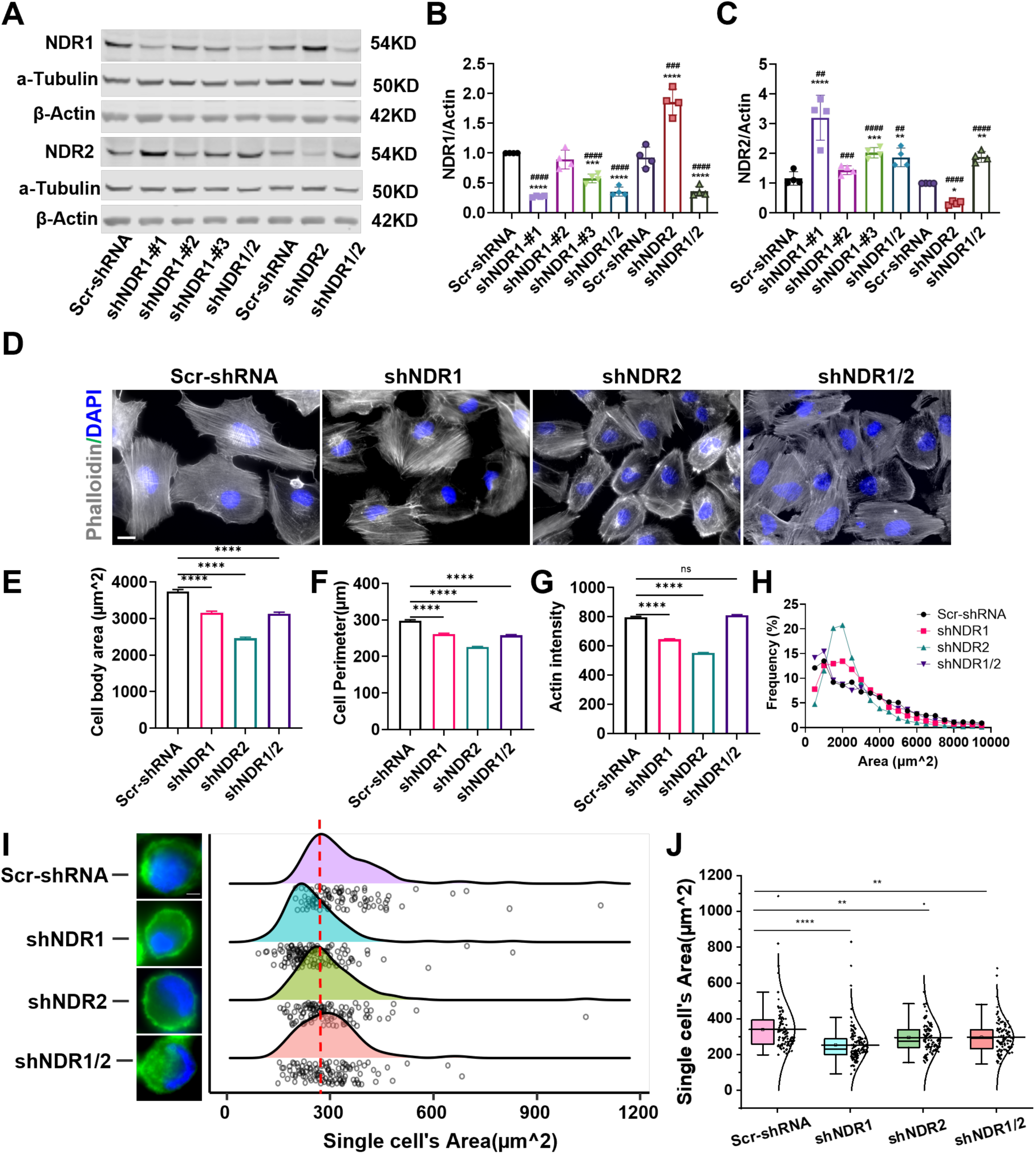
NDR1 and/or NDR2 knock down changes the cell size, shape and actin-cytoskeleton. **(A)** Decreased protein expression level of NDR1 and NDR2 in stably expressed shRNA mediated knock-down human fibroblasts was confirmed by western blot. shRNAs with minimal compensatory effects on NDR2 were selected for further analysis. shNDR1#3 was used for single NDR1 knockdown, while shNDR1#1 was combined with *Ndr2* shRNA to generate the double-knockdown condition (shNDR1/2). ****P < 0.0001, ***P < 0.001, **P <0 .01, *P < 0.05, One-Way ANOVA followed by Bonferroni’s test vs. Scr-shRNA; ####P < 0.0001, ### P< 0.001, ##P < 0.01, compared with Scr-shRNA group, Student’s T-test, n = 4. **(D-G)** Immunofluorescence analysis by Phalloidin staining of fibroblasts followed by an unbiased Opera Phenix Plus High-Content Screening System, which shows a decreased cell size (E-F), and actin intensity (G), 3 days after cell seeding on Ploy-D-lysine coated coverslips; ****P < 0.0001, One-Way ANOVA followed by Bonferroni’s test vs. Scr-shRNA; n = 4902-8457, Bar = 50µm. **(H)** The frequency distribution of cell spreading area. (I-J) Cells were trypsinized and stained with phalloidin (green) and DAPI (blue) for visualization in the mountain map (I). Quantification of cell area is shown in (J).****P < 0.0001, **P < 0.01, One-Way ANOVA followed by Bonferroni’s test vs. Scr-shRNA, n = 96-130.

Together, these observations reveal that NDR kinases play a role in regulating cell morphology, and cytoskeletal organization. Among the three principal types of cytoskeletal filaments, the actin cytoskeleton is the primary force-generating machinery within the cell body (Svitkina, 2018). Actin cytoskeleton–derived forces underlie a wide spectrum of cellular motility processes. Through coordinated polymerization of multiple filaments, the actin cytoskeleton produces protrusive forces, while sliding interactions with bipolar myosin II filaments generate contractile forces. Directed cell motility requires a proper coordination of actin cytoskeleton at the front and rear of the cell (Pontes et al., 2017). Thus, we next asked whether NDR kinases are involved in the regulation of cell motility in fibroblasts.

### NDR kinases are critical for maintaining persistent, directed migration of fibroblasts

We found that NDR knockdown cells exhibited significant alterations in morphology during migration (Fig. 2 A). While control group cells migrated directionally into the open space, NDR knockdown cells displayed more random migration patterns. These findings suggest that NDR kinases may regulate directional cell migration.

**Figure 2.**
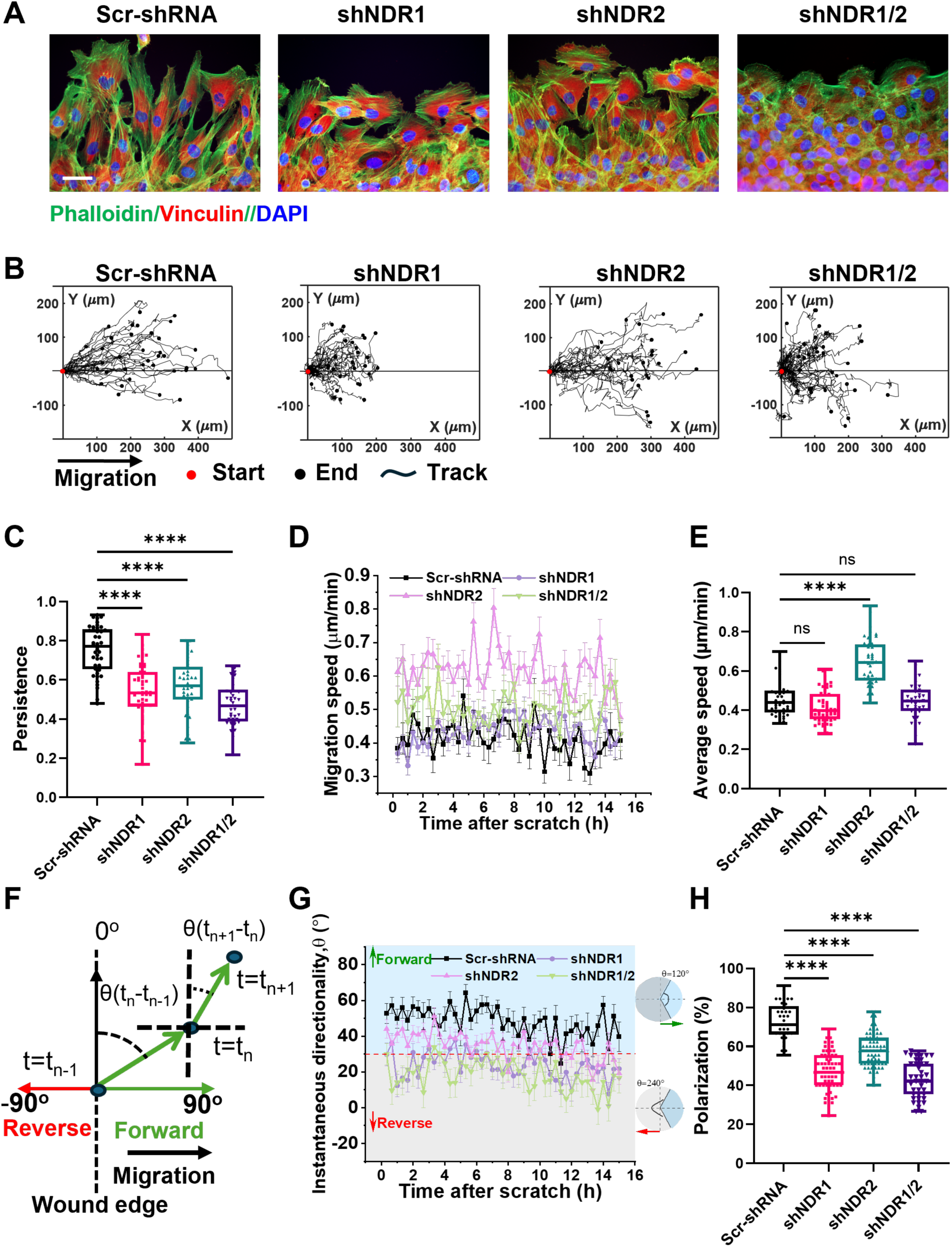
NDR kinases enable direct and persistent cell migration of human fibroblasts. Cells were imaged at 20-min intervals over 15 h using the Incucyte® live-cell imaging system to track individual cell migration. **(A)** Cell monolayers were scratched, cultured for 10 h, and stained with phalloidin (Green) and vinculin (Red) for immunofluorescence visualization, Bar = 50µm. **(B)** Representative trajectories of leading (defined as the first layer of cells at the wound edge) fibroblasts stably expressing shRNA against *Ndr1* and/or *Ndr2* in an *in vitro* wound-healing assay for 15 hours. **(C)** Cell motile persistence, defined as end-to-end displacement divided by total displacement. ****P < 0.0001, One-Way ANOVA followed by Bonferroni’s test vs. Scr-shRNA; n = 38-53. **(D)** Instantaneous cell migration velocity of leading cells. **(E)** Quantified average migration speed per cell over 15 hours. ****P < 0.0001, One-Way ANOVA followed by Bonferroni’s test vs. Scr-shRNA; n = 38-53. **(F)** Schematic of instantaneous cell move directionality. The angle of θ(t_n_-t_n-1_) represents the movement direction between the previous position (t_n-_1) and the current position (t_n_) relative to the wound edge. θ > 0° indicates the cell move towards the wound edge, whereas θ < 0° indicate cell move away from the wound edge. **(G)** Quantification of the instantaneous cell directionality, with θ > 30° defined as polarized cells. **(H)** Quantification of overall average cell polarization over 15 hours, . ****P < 0.0001, One-Way ANOVA followed by Bonferroni’s test vs. Scr-shRNA; n = 37-68.

To evaluate whether NDR kinases influence cell motility in response to release of contact-inhibition, single cell migration behavior was tracked over 15 hours in a monolayer wound healing assay (Mason et al., 2019). Control and NDR kinases knockdown cells were tested in a scratch-induced migration assay, a well-characterized cell model system, and cell motility, including single-cell migration speed, instantaneous directionality of cell movement, square/net displacement, and wound closure rate were evaluated. We found that the NDR1 and/or NDR2 depleted cells moved more randomly and covered shorter distances (shNDR1) than control or shNDR2 cells (Fig. 2 B).

Further, we measured the instantaneous migration speed of single cells at the wound front, designated as leading-edge cells. No significant differences in cell migration speed were observed between control, shNDR1, and shNDR1/2 groups. Conversely, cells with NDR2 knockdown exhibited a marked increase in migration speed (Fig. 2, D-E).

Directional cell migration is indispensable for physiological processes, and pathological conditions such as wound repair require not only cell motility but also tightly regulated directional migration (SenGupta et al., 2021). Thus, we quantified the instantaneous directionality of cell movement. Cell directionality was defined as the angle θ(t_n_–t_n-1_), representing the movement direction of a cell between its previous position (t_n-1_) and current position (t_n_), measured relative to the wound edge at each 20- min interval (Fig. 2 F). All groups of the cells in the leading edge migrated forward with an angle θ > 0° (Fig. 2 G). The NDR kinases knockdown cells display lower instantaneous directionality and reduced persistence than control cells (Fig. 2, G, C).

Furthermore, we found that loss of NDR kinases leads to decreased average cell polarization, where angle θ > 30° defined cells as polarized (Fig. 2 H and Fig. S1 A). Analysis of individual cell motility revealed that mean square displacement and net displacement were reduced in NDR1- and NDR1/2-depleted cells, whereas NDR2 depletion led to increased displacement compared with control cells (Fig. S1, B-C). We also tracked and analyzed the cells seeded at low densities (randomly distributed) and found no significant differences between NDR knockdown and control groups in either average migration speed or square/net displacement (Fig. S1, E-G).

In addition to single-cell motility, we assessed the overall wound closure ratio over 20 hours at 4-hour intervals. NDR1- and NDR1/2-knockdown cells exhibited a significantly reduced wound closure rate, whereas NDR2 depletion showed no significant difference but displayed a trend toward reduced closure (Fig. S1, H, I).

Conversely, NDR1 overexpression enhanced wound closure, resulting in an increased closure ratio compared with control cells (Fig. S1 J). A similar result was observed with a stably Tet-On–inducible cell line in which shRNAs targeting *Ndr1* or *Ndr2* were driven by the H1-2O2 promoter under the control of a TetR element (Fig. S1 K, (Wiederschain et al., 2009)). Wound healing assay results indicated that doxycycline-induced knockdown of either NDR1 or NDR2 led to a reduced wound closure rate (Fig. S1, L– M). Those data further support the point that NDR kinases are key regulators of cell polarity and directed migration in fibroblasts.

### NDR kinases are required for the directional persistence of cell movement

To further assess the mechanisms involved in the control of cell directed migration and migratory cell polarity by NDR kinases, we looked at the Golgi polarization during wound induced migration. The Golgi polarization was defined as the orientation of the Golgi apparatus within ±60° towards the wound edge (Fig. 3 A, (Mason et al., 2019; Nakamura et al., 1995)). Following scratch wounding of fibroblast monolayers, we evaluated Golgi apparatus polarization at the migratory front. Depletion of NDR kinases did not affect Golgi polarization in leading cells, except in the case of NDR2 knockdown (Fig. 3 B-D). In contrast, NDR kinase–depleted cells exhibited reduced polarization compared with control cells in the trailing region (Fig. 3B–D). Overall, ∼50–80% of cells at the wound edge displayed polarized Golgi under both leading and trailing conditions after scratch induction, a frequency significantly higher than that observed in randomly distributed cells (∼33%) (Fig. 3 B–D; and Fig. S2 A–C). More specifically, Golgi angle distribution was similar between control and NDR knockdown cells under leading conditions, whereas in trailing conditions NDR-depleted cells showed a reduced proportion of Golgi within 0–30° and an increased proportion within 90–180° (Fig. 3 E– F).

**Figure 3.**
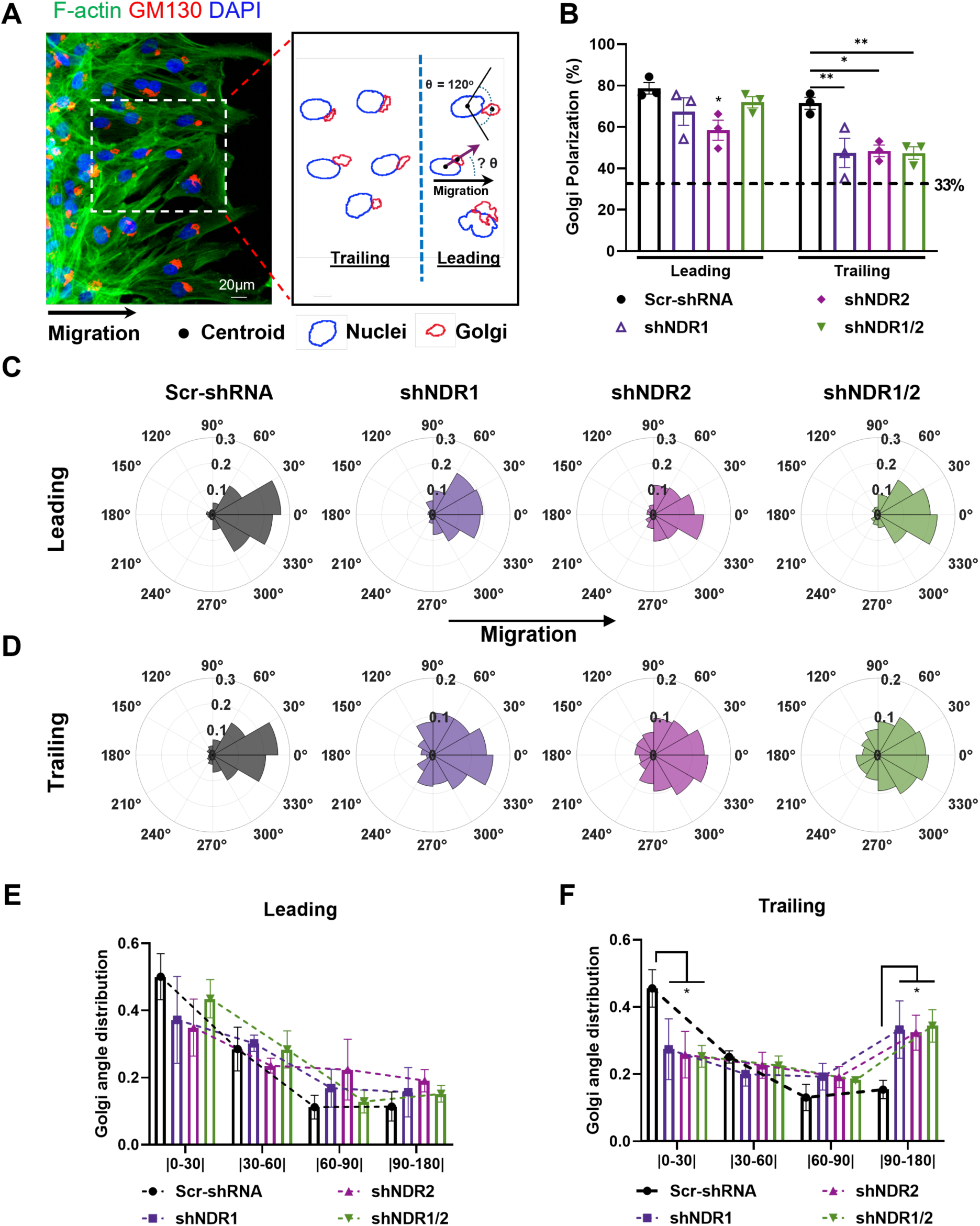
NDR kinases are non-essential for overall Golgi apparatus polarization but required for directional cell migration. **(A)** Immunofluorescence images of fibroblasts fixed 10 h after wound induction. F-actin, Golgi apparatus, and nuclei were visualized with phalloidin (green, Alexa Fluor 488), GM130 (red, Alexa Fluor 647), and DAPI (blue), respectively. A schematic of Golgi polarization measurement shows: blue circles representing nuclei and red circles representing Golgi. The angle (θ) between the wound-edge direction vector and the vector from the nucleus center to the Golgi center was calculated (right panel). Golgi apparatus was considered polarized when θ was within ±60° toward the wound. Leading cells were defined as the first layer of cells at the wound edge, whereas all other cells were classified as trailing cells. Bar = 20µm. **(B)** Quantification of the percentage of Golgi apparatus polarized to the wound in leading and trailing cells. **P < 0.01, *P < 0.05; Two-Way ANOVA followed by Tukey’s post hoc tests. n = 387 - 445 leading cells; n = 647 - 1099 trailing cells with 3 independent experiments. **(C-D)** Rose plot of Golgi apparatus polarization in Scr-shRNA (Dark), shNDR1(Medium purple), shNDR2 (Vibrant purple) and shNDR1/2 (Green) fibroblasts for leading (C) and trailing (D) cells. **(E-F)** Distribution of Golgi apparatus orientation angles to the wound edge in leading (E) and trailing (F) groups, *P < 0.05, One-Way ANOVA followed by Bonferroni’s test vs. Scr-shRNA; n = 3.

These data suggest that NDR kinases are essential for maintaining directional persistence, whereas they exert only a limited effect on directional sensing and polarization during migration initiation.

### NDR kinases spatially and temporally control the dynamics of Cdc42 GTPase

We next investigated the molecular mechanisms underlying the functional specificity of cell migration behavior. Cell migration involves dynamic and spatially regulated changes of actin filament-based structures, including lamellipodia, filopodia, stress fibers, and blebs of membrane as well as the cell-cell/extracellular matrix adhesions (Lawson and Ridley, 2018; Ridley, 2001). For all those cellular events, Rho family small guanosine triphosphatases (GTPases) play a central role in coordinating the cellular responses required for cell migration (Ridley, 2015). To determine whether the conserved NDR kinases are involved in controlling Rho family GTPases activity, we employed GST-CRIB (PAK_23-137_, a Cdc42- and Rac-interactive binding motif that selectively binds GTP-bound Cdc42 or Rac1) and GST-RBD (Rhotekin, which binds GTP-bound RhoA) pull-down assays (Fig. 4 A, and Fig. S3 A). GST pull-down assays revealed increased Rac1 GTPase activity and reduced RhoA GTPase activity in human fibroblasts upon loss of NDR kinases, consistent with previous reports in other species (Fig. S3 B-E, (Emoto et al., 2004; Tang et al., 2015)).

**Figure 4.**
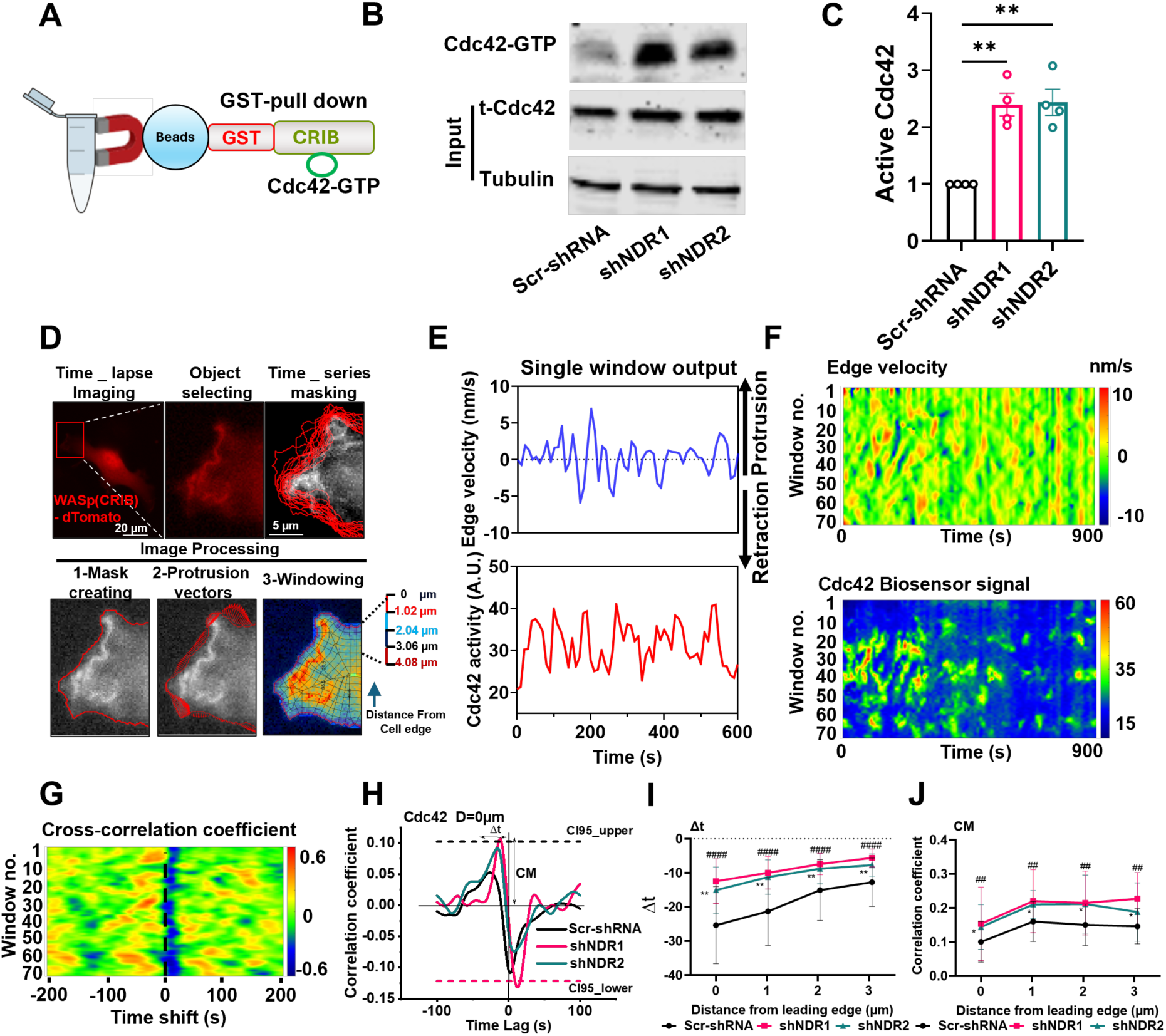
NDR kinases spatially and temporally control the dynamics of Cdc42 GTPase. **(A)** Schematic of the mechanism of GST-pull down assay to analysis the GTP- bounded Cdc42 GTPase. **(B-C)** Western blot analysis the active Cdc42 by GST-Pull down assay. **P < 0.01, One-Way ANOVA followed by Bonferroni’s test vs. Scr-shRNA; n = 4. **(D)** Image analysis pipeline to acquire time series edge dynamics and Cdc42 biosensor (CRIB) activity. Fibroblasts with shRNA-mediated knockdown of NDR1 or NDR2 were stably transduced with a localization-based Cdc42 biosensor (WASP– CRIB–dTomato) and enriched by fluorescence-activated cell sorting (FACS). Time-lapse images were acquired at 10-s intervals for 100 frames. Image regions were segmented according to cell edges, followed by masking, generation of protrusion vectors, and windowing. Within each window, Cdc42 GTPase activity was measured over time. At Distance = 0, a time course of the edge velocity was simultaneously recorded. Bar = 20 µm. **(E)** Time courses of cell edge velocity (blue) and Cdc42 GTPase activity (red) in a representative sampling window. **(F)** The time-course activity map of edge movement and Cdc42 activity (color indicates the level of velocity and GTPase activity). Velocity values along the cell edge were sampled at each time point and sequentially placed into the corresponding columns of the map, generating a spatiotemporal representation of edge movement (top graph). Red represents the protrusion, and blue represents the retraction. Activity maps of Cdc42 GTPase (bottom graph). The activity is displayed as color intensity within a sampling window at Distance = 1.02 – 2.04 µm. **(G)** Cross-correlation coefficients between edge velocity and Cdc42 activity plotted as a function of sampling window position and temporal shift. **(I-J)** Quantification of magnitude of the cross-correlation maximum (CM) and the time lag of cross-correlation maxima as a function of distance D. ####P < 0.0001, ##P < 0.01, **P < 0.01, *P < 0.05; Two-Way ANOVA followed by Tukey’s post hoc tests. n = 11 cells for Scr-shRNA; n = 14 cells for shNDR1 and n = 8 cells for shNDR2 group.

Biochemical evidence suggests that Rac1 and RhoA are mutually inhibitory, and that Cdc42 can activate Rac1 (Arthur and Burridge, 2001; Nimnual et al., 2003; Nobes and Hall, 1995; Ohta et al., 2006; Rottner et al., 1999). Intriguingly, our GST-Pull down assay showed a dramatic increase in Cdc42 activity upon loss of NDR kinases (Fig. 4 B-C), which has yet to be reported in mammalian cells. Cdc42 GTPase plays a central role in regulating cell polarity during asymmetric division, differentiation, and migration in organisms ranging from yeast to mammals (Das et al., 2012; Etienne-Manneville, 2004; Pino et al., 2021).

To investigate the spatiotemporal dynamics of Cdc42 GTPase activity in fibroblasts, we utilized a localization-based biosensor (WASP-CRIB-dTomato) stably expressed in cells with the NDR1 or NDR2 knockdown background (Mahlandt et al., 2023). Time-lapse fluorescence imaging was conducted at 10-second intervals over a 1000-second duration in low density cells. Using a custom MATLAB-based analysis pipeline, we segmented the cell edge and applied automated windowing to define 30 to 100 spatial sampling regions along the cell boundary (Fig. 4 D, analyzed as described in (Lee et al., 2015; Machacek et al., 2009; Noh et al., 2022)). The window width varied according to the cell boundary curvature, while the depth was maintained at a constant 1.02 µm. We tracked the position of the cell edge to identify the status of protrusion and retraction in 300nm-long sectors and calculated edge velocity of each sector over time. These measurements were then compared with the corresponding Cdc42 biosensor signal to assess the temporal coordination between membrane dynamics and GTPase activity, using independently sampled values from adjacent windows. Consistent with previous findings in mouse embryonic fibroblasts (MEFs), we observed oscillatory Cdc42 dynamics at the membrane-a conserved behavior of Rho GTPases and a product of positive and negative feedback mechanisms (Fig. 4 E, (Machacek et al., 2009)). Changes in edge velocity closely mirrored fluctuations in Cdc42 GTPase activity, with a measurable time lag, indicating a temporal correlation between membrane dynamics and signaling activity.

We then extracted locally averaged Cdc42 activity and edge dynamics and visualized them as space-time matrices to capture the spatiotemporal coordination between membrane behavior and Cdc42 GTPase signaling (Fig. 4 F, and Fig. S3 F-H). Membrane dynamics and Cdc42 GTPase activity at the leading edge of the cell both exhibited a similar periodicity of approximately 100 seconds, according to auto- correlation function analysis (see Fig. S3 L-O). We found no significant difference in periodicity of membrane dynamics or Cdc42 GTPase between the Control and NDR kinase knockdown groups (Fig. S3 L-O).

Next, to quantify the relationship between cell membrane dynamics and Cdc42 GTPase activation, we computed the cross-correlation between edge velocity and Cdc42 GTPase activity in each window as previously described (Machacek et al., 2009). High correlation coefficients indicated that GTPase activation was tightly linked to cell protrusion or retraction, whereas lower values showed a less direct relationship. If peak correlation was found at a nonzero time point, it revealed how signaling events and membrane movements were coordinated over time. We observed strong correlations mainly at negative time lags (Fig. 4G, and Fig. S3 F-H (Bottom)), suggesting that Cdc42 GTPase activation lags behind cell edge movement. This finding aligns with earlier results reported in mouse fibroblasts (Machacek et al., 2009). We averaged correlation coefficients across sampling windows for each cell, pooled data from all cells, and fitted a spline to find the significant peak. This quantified the average time lag where membrane dynamics and GTPase activation were most strongly coupled. Our results show that in the control group, Cdc42 activation at the cell edge lags behind membrane protrusion by about 25 seconds, whereas NDR1 or NDR2 knockdown reduces this delay to 12–15 seconds (Fig. 4 H-I). In addition to assessing Cdc42 activation at the cell edge (D = 0 -1.02 µm), we examined Cdc42 activity at varying distances from the cell boundary to obtain comprehensive spatial dynamics information. Correlation analysis was conducted, with calculations of both the maximum correlation coefficients (CM) and their associated time lags (Δt) at different distances from the cell edge. The highest correlation coefficients were found at about 1.02 to 2.04 µm in both the control and NDR kinase knockdown groups, and similar patterns in time lag trends at the cell edge appeared across treatment groups, but with a reduced delay from the cell edge to centra regions (Fig. 4 I-J, and Fig. S3 I-K). The NDR kinase knockdown group demonstrates an increased correlation between Cdc42 activation and cell edge dynamics as compared to the control group (Fig. 4 J).

Taken together, these findings indicate that NDR kinases play a role in controlling cytoskeleton remodeling by regulating, at least in part, the dynamics of Cdc42 GTPase in human fibroblasts. In this manner, NDR kinases, by regulating Cdc42 dynamics and distribution, have a crucial role in controlling exploratory cell behavior and directional cell movement.

### NDR kinases regulate Pard3 protein levels and subcellular localization at cell–cell junctions

Reduction levels of NDR kinases results in hyperactivation of Cdc42 GTPase and premature Cdc42 activation at the leading edge, key events in controlling cell migration polarity. Therefore, we sought to elucidate how NDR kinases regulate Cdc42 activity. To address this, we conducted a bioinformatics analysis using previously published phosphoproteomics data generated from conditional NDR1/2 double knockout mouse hippocampal tissues (Rosianu et al., 2023). Out of 55 potential substrates for NDR kinases (p < 0.05), two were identified as being associated with Rho GTPase signaling: Arhgef11, a guanine nucleotide exchange factor for RhoA, and Pard3, a polarity protein known to act as a mutual antagonist of Cdc42 GTPase (Fig. S4 A-B).

Pard3, together with Par-6 and aPKC, is a component of polarity protein Par complexes, which plays a conserved role in establishing polarity in diverse organisms (Hapak et al., 2018; Nishimura et al., 2005; Zhang and Wei, 2022). The membrane-bound activated Cdc42 directly interacts with the semi-CRIB domain of Par-6, facilitating the recruitment of aPKC. This interaction results in the displacement of Pard3, leading to the formation of Cdc42-Par-6-aPKC complex (Deutz et al., 2025; Vargas and Prehoda, 2023). Pard3 functions as a scaffold protein that facilitates the recruitment of junctional proteins, like Zo-1 (zonula occludens-1) and Zo-2, to their proper location at tight junctions, which is necessary for the assembly and maintenance of functional junctions in epithelia cells (Chen et al., 2017; Zen et al., 2009).

In this study, we tested Pard3 localization and found that NDR kinases knockdown leads to a dramatic mis-location of Pard3 at the cell-cell junctions in both densely grown and scratched fibroblasts monolayers, as shown in immunofluorescence staining (Fig. 5 A-C). Further, when we tested Pard3 protein level by western blot, we found that Pard3 protein level decreased around 50% after NDR1 and/or NDR2 depletion (Fig. 5 D-E). To investigate the location of Pard3 at cell-cell junctions, we performed co-immunostaining for Pard3 and ZO-1. Consistent with our expectations, depletion of NDR1/2 resulted in mis-localization of both Pard3 and ZO-1 protein. In the Scramble shRNA control group, Pard3 and ZO-1 were co-localized at cell-cell junctions, whereas in NDR kinase-depleted cells, both proteins were dispersed and failed to co- localize at cell-cell junctions (Fig. 5 F). To further investigate the impact of NDR1/2 depletion on Pard3 localization, we generated NDR1/2 knockdown fibroblasts stably expressing exogenous Pard3 fused to GFP at the C-terminus (Fig. 5 G). Live-cell imaging revealed pronounced disruption of Pard3-GFP localization at cell-cell junctions in both randomly distributed and scratch-wounded cell cultures of NDR1/2 knockdown cells. Consistent with these observations, fixed-cell imaging confirmed that the mis- localization of Pard3-GFP persisted under both conditions (Fig. 5 H). Moreover, a confocal microscopy analysis and 3-D reconstruction demonstrated that Pard3 forms clusters throughout the cell body following NDR kinase knockdown (Fig. S4 L). To validate that the observed changes in Pard3 protein were a direct consequence of NDR kinase depletion, we established stable Tet-on inducible cell lines that express shRNA targeting to *Ndr1* or *Ndr2* (Wiederschain et al., 2009). Upon doxycycline induction, shRNA-mediated knockdown of either Ndr1 or Ndr2 resulted in a reduction of Pard3 protein levels, as shown in (Fig. S4 C–J). Notably, both Pard3 and the junctional protein Zo-1 exhibited pronounced mis-localization in the doxycycline-treated Ndr1 and Ndr2 Tet-on shRNA groups. In contrast, the Scramble control group displayed no alterations in Pard3 or Zo-1 localization, regardless of doxycycline exposure (Fig. S4 K).

**Figure 5.**
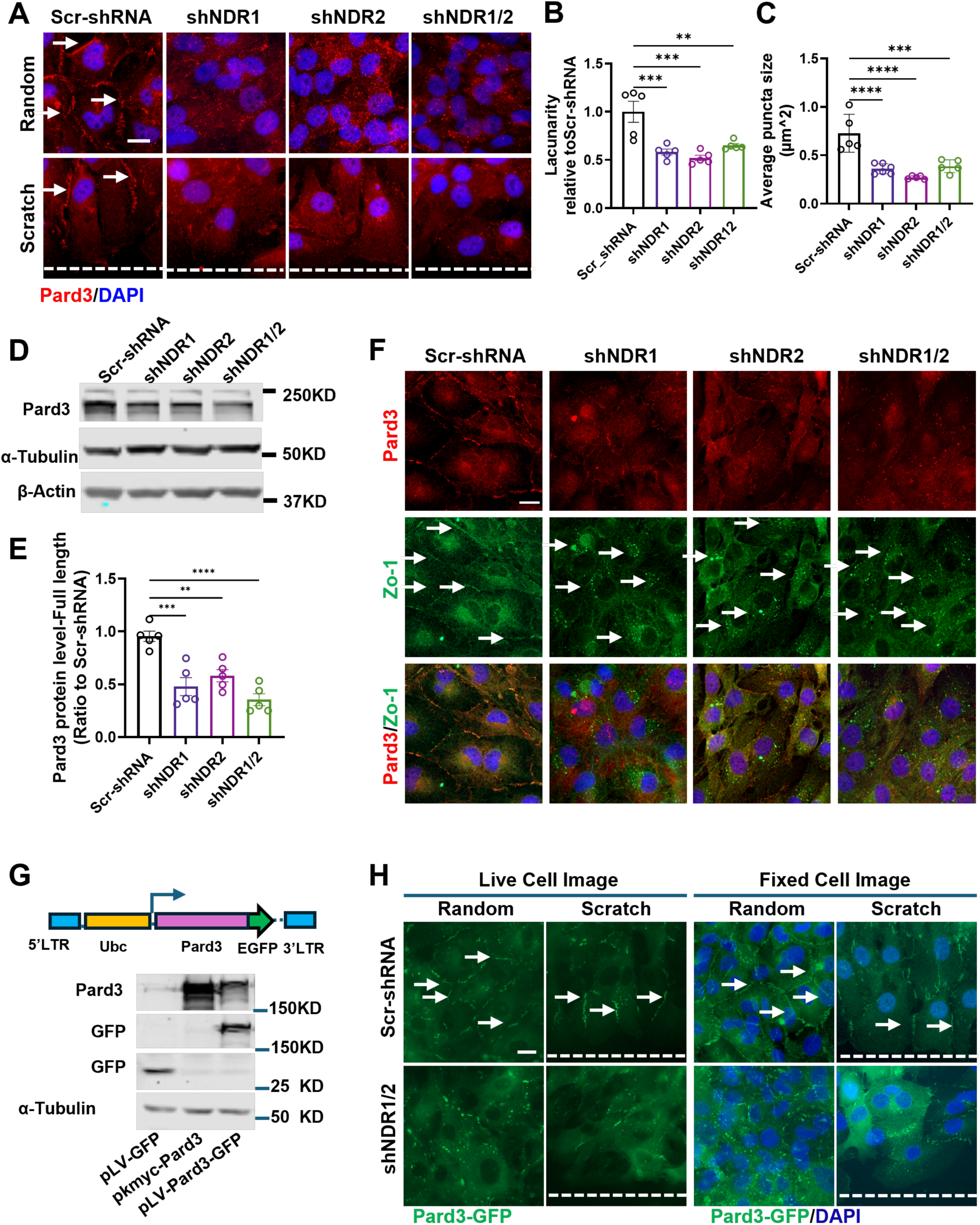
NDR kinases regulate Pard3 expression and subcellular localization at cell–cell junctions. **A-C** Fibroblasts were fixed 10 h after wound scratch and subjected to immunofluorescence analysis. Pard3 was visualized with Alexa Fluor 647 (A). Junctional organization was quantified by lacunarity (B) and average puncta size (C)., ****P < 0. 0001, ***P < 0.001, ** P < 0.01, One-Way ANOVA followed by Bonferroni’s test vs. Scr-shRNA; n = 4. **(D-E)** Pard3 protein expression levels were analyzed by western blot, ***P < 0.001, ** P < 0.01, One-Way ANOVA followed by Bonferroni’s test vs. Scr-shRNA; n = 5. **(F)** Co-localization of Pard3 and the junctional protein ZO-1 was assessed by co-immunofluorescence staining. Pard3 was visualized with Alexa Fluor 647 (red), ZO-1 with Alexa Fluor 488 (green), and nuclei with DAPI(Blue). **(G)** Schematic of the pLV-Pard3-GFP lentiviral construct driven by the Ubc promoter. Construct integrity was confirmed by sequencing and western blot analysis. **(H)** Fibroblasts with shRNA-mediated knockdown of NDR1/2 were infected with Pard3- GFP–expressing lentivirus and enriched by FACS. Pard3 localization was visualized by GFP in both live-cell and fixed-cell imaging. The white dashed line indicates the scratch wound edge. Scale bars in (A, F, H): 20 µm.

In summary, our findings demonstrate that NDR kinases are essential regulators of the polarity protein Pard3, influencing both its expression protein levels and subcellular localization. The depletion of NDR kinases leads to mis-localization and reduced levels of Pard3. Since Pard3 may affect active Cdc42 GTPase pools (Vargas and Prehoda, 2023), this disruption in the Pard3 and Cdc42 signaling may ultimately impair cell polarity and migration persistency in fibroblasts.

### NDR kinases phosphorylate Pard3 at Ser144, and the S144A mutation abolishes the rescue effect of Pard3-WT in wound healing

To investigate whether NDR kinases directly phosphorylate Pard3 at the predicted consensus motif (H.R..[S/T]) (Rosianu et al., 2023), specifically at serine 144 (S144), a site that was identified by phosphoproteomic analysis (Rosianu et al., 2023), we generated a truncated N-terminal fragment of Pard3 containing the S144 site as substrate, which was expressed with a C-terminal His tag in E. coli DE3 cells (Fig. 6A, and Fig. S5 C-D). To obtain activated NDR kinases, both wild-type (WT) and kinase-dead (KD, K118A for NDR1 and K119A for NDR2 kinase) variants of NDR1 and NDR2, each with an N-terminal His tag, were expressed in HEK293T cells under the control of a CMV promoter (Fig. S5 A-B). The expressed constructs were all verified by sequencing and western blot analysis. Purified NDR kinase proteins and the corresponding N-terminal Pard3 substrates (WT/S144A mutants) were then applied in an *in vitro* thiophosphorylation kinase assay to assess phosphorylation activity (Fig. 6 C–F). Notably, substituting serine 144 with alanine in the Pard3 fragment, or introducing kinase-dead mutations (K118A/K119A) in NDR1/2 kinases, led to a marked decrease in the thiophosphate ester signal (Fig. 6 C-F), indicating a loss of phosphorylation activity by those mutations.

**Figure 6.**
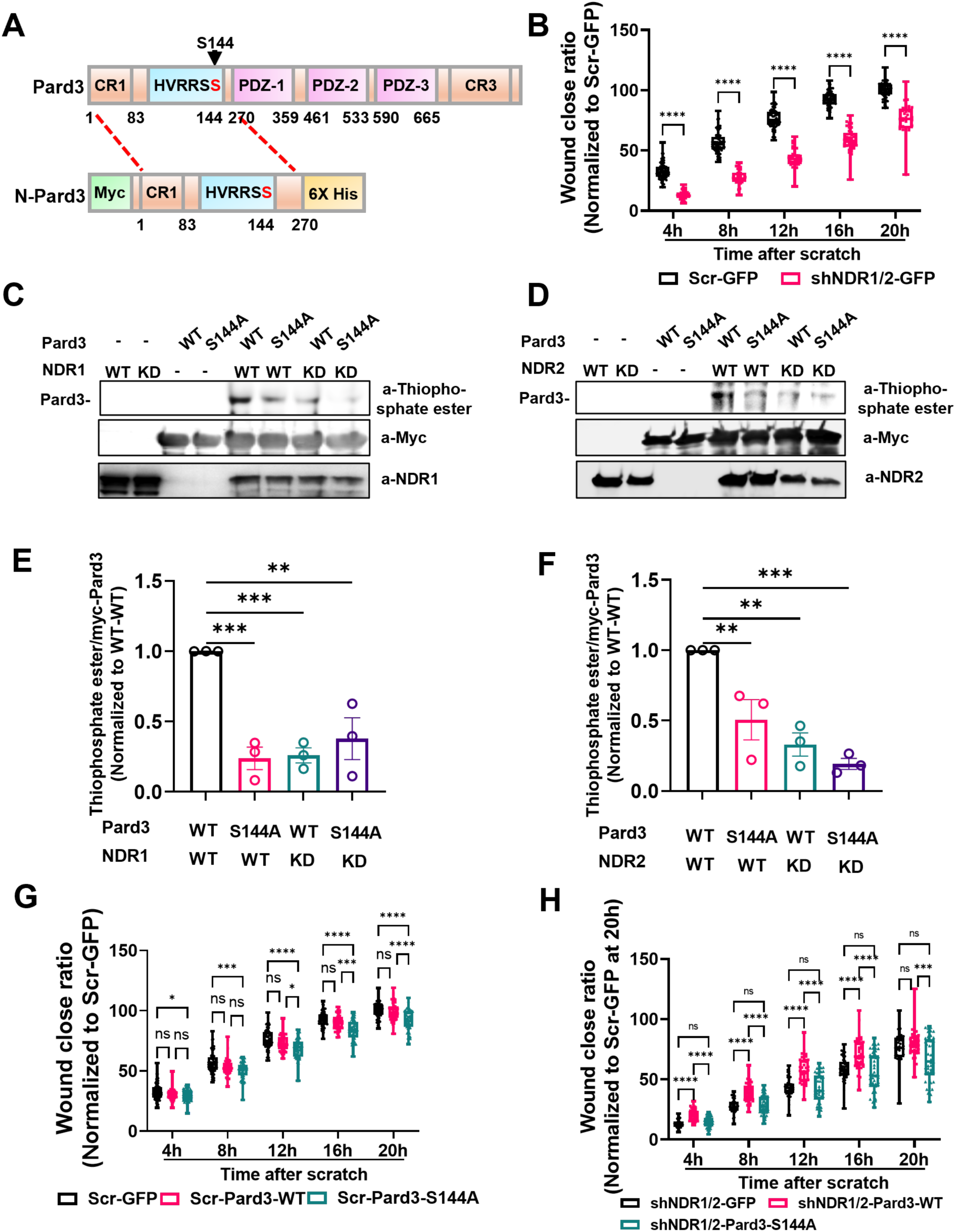
NDR kinases phosphorylate Pard3 at Ser144, and the S144A mutation abolishes the rescue effect of Pard3-WT in wound healing. **(A)** Schematic of Pard3 (top) highlighting the consensus NDR kinase phosphorylation motif (H.R..[S/T]); The N-terminal region of Pard3 (tagged with Myc at the N-terminus and 6×His at the C-terminus) contains the consensus phosphorylation site at Ser144. Wild-type (WT) and S144A mutant constructs were cloned into the pET22b backbone for inducible expression in E. coli DE3 cells (bottom). **(B)** Fibroblasts stably expressing lentivirus encoding GFP alone were subjected to wound-healing assays as controls; ****P < 0.0001; Two-Way ANOVA followed by Tukey’s post hoc tests, n = 40. **(C-F)** Validation of Pard3 phosphorylation at Ser144 by in vitro kinase assays. NDR1 and NDR2 (WT or kinase-dead [KD], , K118A for NDR1 and K119A for NDR2) were purified from HEK293T cells and incubated with purified WT N-Pard3 or S144A-mutant N-Pard3 protein. Reactions were performed in the presence of ATP-γ-S, and thiophosphorylation was detected by western blot using an anti–thiophosphate ester antibody, ***P < 0.001, **P < 0.01; One-Way ANOVA followed by Dunnett’s post hoc tests, n = 3. **(G-H)** Rescue wound-healing assays were performed to evaluate the effect of exogenous Pard3 or the Pard3-S144A mutant on fibroblast migration following NDR1/2 knockdown over 20 h, ****P < 0.0001, ***P < 0.001, *P < 0.05; Two-Way ANOVA followed by Tukey’s post hoc tests, n = 40-46.

To clarify the function of Pard3-S144 in cell migration, we generated human fibroblasts stably expressing either exogenous wild type Pard3 fused to GFP (Pard3- WT-GFP) or a mutant form where serine 144 was replaced with alanine (Pard3-S144A- GFP). Those constructs were introduced into both control (Scramble shRNA) and NDR1/2 knockdown backgrounds. Control experiment confirmed that GFP expression alone does not affect the reduced wound closure ratio seen with NDR1/2 knockdown, as measured by an Incucyte live-cell imaging (Fig.6 B). In the scramble shRNA group, expression of wild type Pard3 (Pard3-WT-GFP) did not alter wound closure efficiency. In contrast, cells expressing Pard3-S144A-GFP exhibited a small but significant decrease in wound closure efficiency (Fig. 6 G). Importantly, in NDR1/2 knockdown cells, Pard3- WT-GFP partially rescue the impaired wound healing phenotype, whereas Pard3- S144A-GFP failed to do so, completely abolishing the rescue effect (Fig. 6 H). These results demonstrate that the phosphorylation of Pard3 at serine 144 is essential for its role in promoting cell migration and wound repair.

### Downregulation of NDR1 diminished re-epithelialization in human ex vivo wound healing and suppresses keratinocyte migration in vitro

Given that NDR kinases have an influence on fibroblasts migration in wound healing assays, we hypothesized that NDR kinases have a role in human skin wound healing.

To evaluate the impact of NDR kinase on re-epithelialization, we employed a human *ex vivo* acute wound model to evaluate wound healing processes. Human abdominal skin tissue with fresh wounds was infected for 6 hours with lentivirus expressing shRNAs against *Ndr1 and Ndr2*. Epithelial tongue length was quantified at day 4 post-wounding. We were successful at knocking down significantly NDR1 in human *ex vivo* skin (Fig. 7 C-D). We found that NDR1 knockdown significantly reduced wound reepithelialization (Fig. 7 A-E). Parallel experiments in HaCaT keratinocytes with inducible NDR1 knockdown corroborated these results, showing dramatically impaired wound closure in vitro. (Fig. 7 F-H, and Fig. S6 A-B). Thus, these findings connect the molecular role of NDR kinases in cytoskeletal dynamics and cell polarity to an important physiological outcome in skin wound healing.

**Figure 7.**
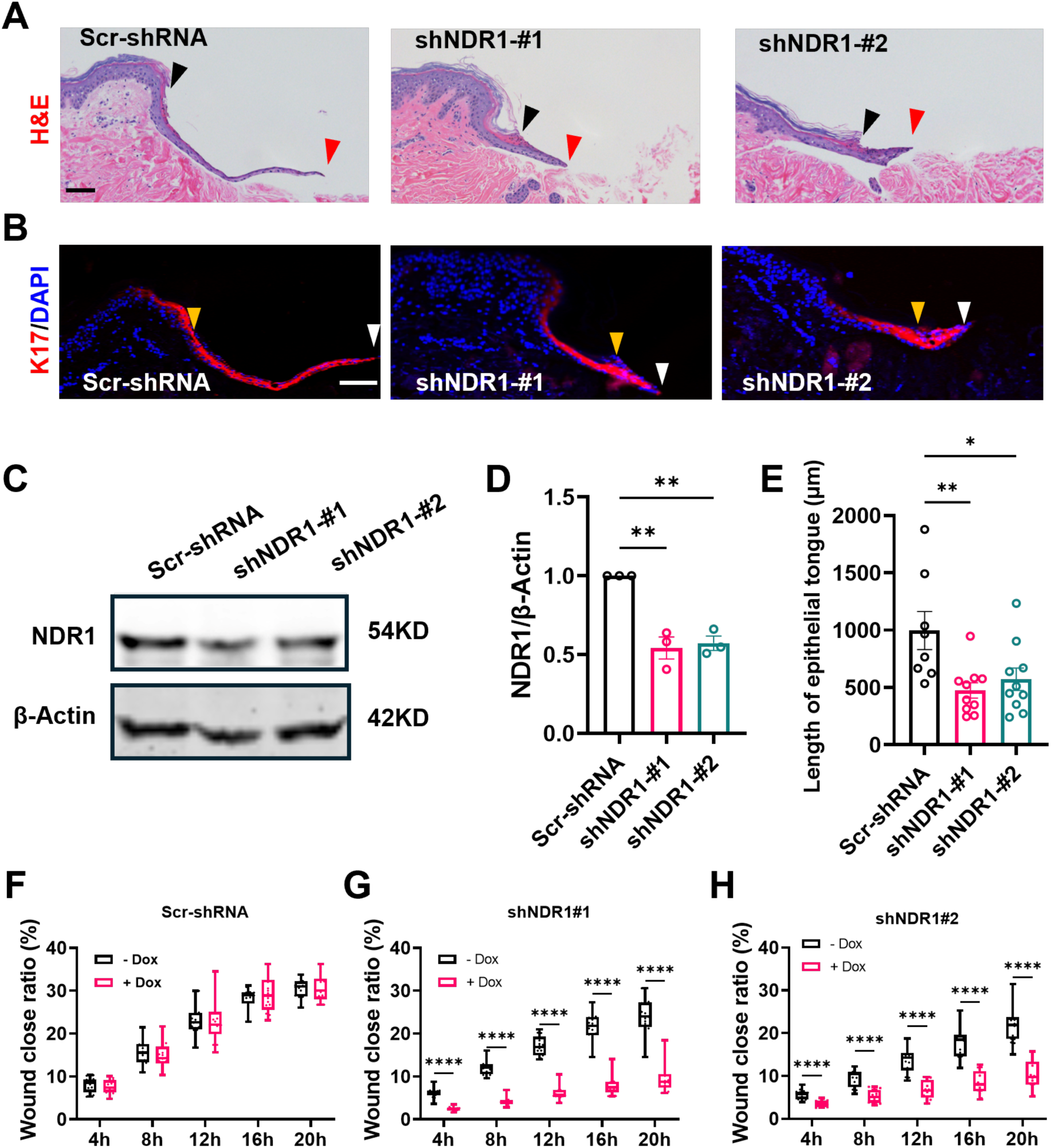
Downregulation of NDR1 diminished re-epithelialization in human ex vivo wound healing and suppresses keratinocyte migration in vitro. **(A)** Hematoxylin and Eosin (H & E) staining of human skin *ex vivo* wounds, treated with lentivirus particles encoding scramble shRNA or shRNA targeting NDR1; black arrowheads indicate the original site of wounding; red arrowheads mark the edge of the migrating epithelial tongue. Bar = 100 µm. **(B)** Immunofluorescence staining of human skin ex vivo wounds with anti–cytokeratin 17 (red) to label activated keratinocytes; nuclei were stained with DAPI (blue); yellow arrowheads indicate the original site of wounding, and the white arrowheads indicate the edge of the migrating epithelial tongue. Bar = 100 µm. **(C-D)** Western blot confirming NDR1 knockdown by lentiviral shRNA. **P <0.01, One-Way ANOVA followed by Bonferroni’s test vs. Scr-shRNA; n = 3. **(E)** Quantification of re-epithelialization by histomorphometric analysis; **P <0.01,*P < 0.05, One-Way ANOVA followed by Bonferroni’s test vs. Scr-shRNA; n =3 biological replicates and 2 to 4 technical replicates per group. **(F-H)** Scratch wound-healing assay of HaCaT keratinocytes with inducible NDR1 shRNA. Cell migration was monitored and analyzed using the Incucyte® live-cell imaging system. ****P < 0.0001; Two-Way ANOVA followed by Tukey’s post hoc tests, n = 16.

## Discussion

While the LATS family of kinases have been widely studied across various species since the *Dbf2* gene was first identified in *S. cerevisiae* (Johnston et al., 1990), the related family of NDR (nuclear dbf2-related) kinases is much less studied. The identification of NDR homologs in *Neurospora crassa* (Yarden et al., 1992), humans, *Drosophila melanogaster*, *C. elegans* (Millward et al., 1995), *S. pombe* (Verde et al., 1995; Verde et al., 1998), and *S. cerevisiae (Geyer et al., 1999)*, enabled initial research into NDR kinase functions. In higher eukaryotes, research on NDR kinase function initially centered on their role in regulating neuronal cell shape (Emoto, 2011; Emoto et al., 2004; Leger et al., 2018; Ultanir et al., 2012). This study uncovers a novel function of NDR kinases in wound healing by promoting sustained fibroblast motility and demonstrates their role in regulating Cdc42 activity in higher eukaryotes. Using RNA interference-based genetic approaches, we showed that Cdc42 and Pard3 act as essential downstream effectors of NDR kinases in regulating cell migration. We found that, first, knockdown of NDR1 and/or NDR2 kinase leads to a dramatic change in cell morphology, including reduced cell size, altered shape, and disrupted actin cytoskeletal organization. Second, cell migration persistence and polarity are impaired after NDR kinases depletion, assessed by cell wound healing assays, single-cell tracking and Golgi polarization assays. Third, the knockdown of NDR1 or NDR2 results in increased activity of Cdc42, a key regulator of cell polarity. Cdc42 biosensor analyses indicate that NDR kinases are capable of spatially and temporally regulating the dynamics of Cdc42 GTPase. Fourth, NDR kinases regulate the protein expression and subcellular location of polarity protein Pard3, ultimately influencing the activity of Cdc42 GTPases. NDR kinases phosphorylate Pard3 at Ser144, and this post-translational modification is essential for Pard3 function in promoting cell migration. Finally, NDR1 knockdown impairs re-epithelialization in a human *ex vivo* wound healing model and suppresses keratinocyte migration *in vitro*. Together, our results have implication in targeting NDR kinases as a potential therapeutic intervention for wound healing.

### Control of cell morphogenesis by NDR kinases

NDR kinases, part of the NDR/LATS subfamily of AGC kinases, are key to cell morphogenesis across species (Hergovich et al., 2006; Sharif and Hergovich, 2018; Tamaskovic et al., 2003). In *S. pombe*, dysfunction of NDR kinase Orb6 induces abnormal cell shape (Verde et al., 1998). In Drosophila, Tricornered (Trc, the counterparts of mammalian NDR1/2) is critical for dendritic tiling of neuron cells (Emoto, 2011; Emoto et al., 2004). In higher eukaryotes, impaired NDR kinases increase proximal dendrite branching in mice neurons (Ultanir et al., 2012). While the importance of NDR kinases on cell morphogenesis has been explored in neurons and *S. pombe*, their contribution to fibroblasts’ morphology has been unclear. Our work reveals that NDR kinases are essential to maintain normal cell shape, size and proper actin assemble in human fibroblasts. We found a reduction in cell spreading area after knocking down NDR1 and/or NDR2 in cells cultured on plastic substrates, and also a reduction in cell volume in suspended fibroblasts detached by trypsin. Unlike the role of NDR kinases in restricting dendritic branching in neurons, our findings indicate that NDR kinases promote cell spreading during fibroblasts growth, which may play a role in fibroblasts activation(Plikus et al., 2021).

Chemical and physical insults induce resting fibroblasts into myofibroblasts through cytoskeleton remodeling, promoting tissue repair (Plikus et al., 2021). Mammals possess three cytoskeletal systems: the actin cytoskeleton, intermediate filaments, and microtubules, all of which must function together for efficient cell migration (Seetharaman and Etienne-Manneville, 2020). Actin dynamics spatially and temporally regulate cell protrusion, adhesion, contraction, and retraction from front to rear (Tang and Gerlach, 2017). NDR1 was shown to facilitate cytoskeleton assembly by interacting with cytoskeleton-related proteins (eg. Actin (ACTB)) during autophagy (Klimek et al., 2019) and regulate sarcomere assembly by modulating Rbm24 protein stability (Liu et al., 2017). In this study, we found an overall decrease of F-actin intensity (Fig. 1 D and G), which suggests a change of actin cytoskeleton remodeling.

### NDR kinases mediate cell migration and polarity

NDR1 and NDR2 share about 87% sequence identity but have different mammalian organ distributions: NDR1 is abundant in immune organs (thymus, spleen, lymph nodes), while NDR2 is highly expressed in the colon and brain, reflecting their distinct functions (Rehberg et al., 2014; Schmitz-Rohmer et al., 2015; Stork et al., 2004). Here, we reported that NDR kinases have distinct effects on cell motility in wound healing assays. NDR2 knockdown significantly increases migration and motility; this effect is reversed by additional NDR1 knockdown (shNDR1/2 group). Knocking down NDR1 alone reduces cell motility (see Fig. 2 D and E, and Fig. S1 B and C). These findings align with earlier reports: naive T cells lacking both NDR genes show a significant reduction in cell motility, whereas naive T cells from NDR2 knockout mice tend to migrate at a higher speed (Tang et al., 2015).

Cell motility, as well as the directionality and persistence of migration, are fundamental for effective cellular movement (Petrie et al., 2009; Seetharaman and Etienne-Manneville, 2020). In this study, we observed that NDR kinases may influence cell migration directionality, initially identified by chance during F-actin staining of scratched fibroblasts (Fig. 2A). The unique arrangement of F-actin filaments observed in the NDR kinase knockdown group indicates that NDR kinases may play a role in controlling cell polarity during migration. Further single-cell real-time behavioral analyses revealed reduced migration persistence and cell polarization following NDR kinase knockdown. Additionally, Golgi polarization assessments in wound-scratched fibroblasts confirmed that NDR kinases are required for maintaining directional persistence; however, they do not appear to be critical for sensing or initiating motile cell polarization. Since this study is limited to two-dimensional cell culture analysis, further research is required to determine how fibroblasts behave in the more complex environments when NDR kinase expression is disrupted, such as a 3-D cell culture system. Additionally, other NDR effectors may participate in the control of motility: for example YAP and TAZ, transcriptional coactivators acting downstream of NDR/LATS kinases, regulate cytoskeletal and focal adhesion maturation to facilitate sustained motility in endothelial colony-forming cells (ECFCs) (Mason et al., 2019). Collectively, these findings demonstrate that NDR kinases are necessary for the establishment of front-rear cell polarity and essential for persistent cell migration.

### NDR kinases regulate the activity of Rho family GTPases

Efficient cell migration depends on the dynamic assembly and breakdown of actin structures like lamellipodia, filopodia, and focal adhesions, which are mainly regulated by the Rho family of small GTPases (RhoGTPases). During two-dimensional cell migration, cells at the leading edge extend lamellipodia and filopodia as a result of actin polymerization advancing the plasma membrane (Lawson and Ridley, 2018). Classic Rho GTPases, including Rac1 and Cdc42, play a key role in regulating lamellipodia and filopodia formation. Rho GTPases are activated through the action of guanine nucleotide exchange factors (GEFs) that stimulate the exchange of a bound GDP nucleotide for GTP, whereas their inactivation is facilitated by GTPase-activating proteins (GAPs) via catalysis of GTP hydrolysis (Bos et al., 2007).

NDR kinases have been proven to play a conserved role in regulating the activity of Rho GTPases. For example, in *S. pombe*, NDR Orb6 controls its substrates, including Cdc42 regulators and other components, which cooperatively contribute to regulate polarized cell growth (Das et al., 2012; Das et al., 2015; Das and Verde, 2013; Das et al., 2009; Doyle et al., 2025; Nunez et al., 2016). NDR Trc negatively regulates Rac activity to control dendritic branching in *Drosophila* neurons, since expression of a kinase-dead mutation of NDR *Trc* leads to a higher Rac1 activity (Emoto et al., 2004). In mice thymocytes, knocking down both NDR1 and NDR2 reduces RhoA activity, which in turn hampers the RhoA-driven polarization of the actin cytoskeleton and ultimately leads to a decreased cell migration (Tang et al., 2015). Consistent with these reports, our data shows that loss of NDR kinases increases Cdc42 and Rac1 activity and decreases RhoA activity in fibroblasts (Fig. 4 B-C; Fig. S3 A-E) highlighting the conservation of these controls across species and across large evolutionary distances.

Interestingly, our findings in fibroblasts differ from those reported in HEK293T cells by Ohta et al., who found that STK38 (NDR1) knockdown reduced Rac1 and Cdc42 activation in HEK293T cells (Ohta et al., 2025). This discrepancy likely reflects cell-type-specific regulation, as fibroblasts and HEK293T cells differ substantially in cytoskeletal organization, signaling context, and differentiation state.

In our hands, Image analysis of the Cdc42 biosensor, in agreement with GST-pull down assay results, demonstrates that NDR kinase regulates Cdc42 activity both spatially and temporally. When NDR kinase is knocked down, Cdc42 is activated earlier, promoting cellular protrusion events, and, since Cdc42 GTPase plays a key role in directing polarized cell growth, altering cell migration.

### NDR kinases regulate Pard3 cellular localization

Cell migration begins by forming a front-rear polarity axis, which is important for directional and cell migration persistence. The polarity protein Pard3, which is one of the NDR target candidates identified by phosphoproteomic analysis (Rosianu et al., 2023), was shown to be regulated by NDR kinases both by phosphorylation and physical interaction in neurons. Yang et.al reported that NDR kinases phosphorylation of Pard3 at Ser383 polarizes the distribution of Pard3 and reinforces axon specification in mice neurons (Yang et al., 2014). We found that Pard3 contains a consensus NDR kinase phosphorylation site at the N-terminal Ser144, near the CR1 domain, which is confirmed by mass spectrometry (Rosianu et al., 2023). Kinases such as PAR1 have been reported to also phosphorylate this site. Once S144 on Pard3 is phosphorylated, 14-3-3 protein binds to Pard3, which then trans-locates from tight junctions to the cytosol, however some mechanistic details are still unclear (Lv et al., 2015; Yang et al., 2012). In particular, Yamashita et al. indicated that phosphorylation of Pard3 at S144 may positively regulate cell tight junction formation, although the exact mechanism remains unknown (Yamashita et al., 2020). On the other hand, Hurd et.al demonstrated that 14- 3-3 interaction with phosphorylated Pard3 at S144 is critical for cell polarity in mammalian epithelia (Hurd et al., 2003). It is possible that phosphorylation of Pard3 at S144 affects Pard3 stabilization through its interaction with 14-3-3 protein. This interaction may, in turn, facilitate the coordinated assembly of Pard3 into cell-cell junctions, acting as a reservoir for the scaffold protein. Nevertheless, this hypothesis requires additional research to be thoroughly evaluated; for instance, 14-3-3 binding to phosphorylated Pard3 at S144 may inhibit Pard3 phase separation into clusters in the cytoplasm or at the membrane (Liu et al., 2020). Additionally, NDR1 and 2 kinases were reported to physically interact with Pard3 both at the N-terminus ( containing the CR1 and PDZ1 domain, where the CR1 domain is critical for self-oligomerization of Pard3) and at the C-terminus (amino acids 1116–1356) (Yang et al., 2014), indicating a specific regulatory manner by which NDR kinase acts on Pard3. NDR kinases have a role in modulating the stability of targets and were reported to stabilize MYC to support the growth of B-cell lymphoma as well as bind to PD-L1 protein to inhibit ubiquitination-mediated PD-L1 degradation. Furthermore, NDR1 physically interacts with BAG3 to mediate autophagy (Bisikirska et al., 2013; Fu et al., 2024; Klimek et al., 2019). Thus, NDR kinase may regulate Pard3 stability both in a kinase-dependent manner and in a kinase-independent manner, by physical interaction.

Our study, detailed in this paper, found that NDR kinase knockdown significantly disrupts Pard3 subcellular localization and reduces Pard3 protein levels in human fibroblasts. At high cell densities, Pard3 localizes to cell-cell junctions whereas it appears mostly diffused at low cell densities. When a scratch in a wound closure assay offers free space with relatively low cell densities, Pard3 moves away from cell-cell junctions in cells located at the leading edge (Fig. 5 A). We found that loss of NDR kinase causes Pard3 protein to delocalize form cell-cell junctions and form clusters throughout the cell body (Fig. 5 A, F and Fig S4 K), suggesting that this alteration may affect cell polarization during migration, and change migration persistence. Pard3 S144 is phosphorylated by NDR kinase *in vitro*, and Pard3 wildtype partially rescues cell migration defects from NDR kinase loss, whereas Pard3 S144A does not. Consistent with the previous report that NDR kinase can phosphorylate Pard3 at S383, mutation of Pard3 S383A also failed to rescue cell migration defects by NDR kinase knockdown (data not shown).

Intriguingly, Pard3 and Cdc42 can form mutually exclusive complexes with the Par-6/aPKC complex in a purified protein qualitative binding assay (Vargas and Prehoda, 2023). We hypothesize that NDR kinases regulate Pard3 localization to the cell-cell junctions and protein expression, which would limit the amount of active Cdc42 at the cell-cell junctions. Our observations suggest that loss of NDR kinase decreases Pard3, which may further result in ectopic activation of Cdc42 GTPase. Elevated Cdc42 would promote filopodia formation, aiding cell migration, but its aberrant activation could disrupt directional sensing and proper cell front-rear polarization. However, the mechanisms by which Cdc42 is activated and how NDR kinase regulates Pard3 expression, and its cellular localization remain unclear, requiring further investigation to be fully understood.

Finally, we found that NDR1 plays a role in human skin *ex vivo* wound healing assays. We further extended these findings using keratinocytes *in vitro* wound healing assays. These results connect the role of NDR kinases in cell motility to a physiological process important to human health. Further research will be needed to fully elucidate the more complex role of NDR kinases and Pard3 in human skin wounds.

In summary, this study identifies NDR1/2 kinases as key regulators of cell polarity and migration in human fibroblasts via Pard3 phosphorylation and Cdc42 activity. These findings enhance our understanding of cell migration mechanisms and in the future, may inform better treatments for wound healing.

## Materials and methods

### Cell culture

Human foreskin fibroblasts (HFF) were gifts from Dr. John M. Murray (Indiana University at Bloomington). HFF cells were grown in 50% high glucose DMEM + 50% Ham’s F10 Nutrient Mix + 10% FBS (Corning, 35-011-CV) supplemented with 1mM sodium pyruvate and 1X Antibiotic-Antimycotic (ThermoFisher, 15240062). The human embryonic kidney cell line (HEK 293T; clone T7) was obtained from the American Type Culture Collection (ATCC), which was cultured in media containing high glucose DMEM with 10% FBS and 1X Antibiotic-Antimycotic. The human keratinocytes cell line HaCaT (AddexBio. T0020001) was gift from Dr. Ivan Jozic, Ph.D (University of Miami). The HaCaT cells were cultured and maintained in DMEM without calcium supplemented with 10% FBS, 1X Antibiotic-Antimycotic and 1X GlutaMAX (Gibco, 35050061). All cells were cultured and maintained in a 5% CO2 incubator at 37℃, unless otherwise indicated.

### Generation of gene knock-down cells with shRNA

Lentiviral vectors that carrying specific shRNA sequences (Scramble as control) were constructed by VectorBuilder. The shRNA was driven by a U6 promoter and marked by TagBFP or mCherry for flow cytometry selection. Three shRNA plasmids targeting *Ndr1* and two shRNA plasmids targeting *Ndr2* were constructed, and the one with less compensation *in vitro* was used *in vivo* human skin experiments. The sequences of human NDR1, NDR2 are: 5’-GTATTAGCCATAGACTCTATT-3’ (shNDR1-#1), 5’- TCGTGTATAGTCATCGAATAT-3’ (shNDR1-#2), 5’-CAGCAAGGGCCATGTGAAACT-3’ (shNDR1-#3), 5’-TCCGAGCAGAAAGAGATATTT-3’ (shNDR2-#1), 5’-GAAGAAGGATTAGCAGATGAA-3’ (shNDR2-#2). The non-targeting scramble shRNA sequence was 5’-CCTAAGGTTAAGTCGCCCTCG-3’ (Scr-shRNA).

The shRNA plasmids combined with Δ8.2 and VSV-G plasmids were co-transfected into HEK293T cells for virus packaging using Lipofectamine 3000 (eg. T25 flask with 6μg of shRNA plasmid, 6μg of Δ8.2 and 1μg of VSV-G plasmid) (ThermoFisher, L3000008).

The supernatant that contains viruses was collected at 24, 48 and 72 hours and filtered through 0.45μm syringe filters (Fisher Sci). The supernatant with 50% fresh DMEM media and 10μg/ml polybrene was added to HFF cells. After infection with lentivirus, HFF cells that express BFP or mCherry fluorescence tag were enriched by fluorescence-activated cell sorting (FACS). After expansion, the cells were verified by western blotting to examine changes in protein levels.

### Molecular cloning and Plasmids

The Pard3-containing plasmid was Addgene # 19388 (deposited by Ian Macara), NDR1- containing plasmid was Addgene # 23811, and NDR2-containing plasmid was Addgene # 23810 (deposited by William Hahn, David Root). pLV-CMV-EGFP -T2A- NDR1-WPRE was created by PCR amplifying NDR1 (primers: FW: 5’- AGAATCCTGGCCCAGGATCCATGGCAATGACAGGCTCAAC-3’, and RV, 5’- ATAAGCTTGATATCGAATTCCTATTTTGCTGCTTTCATGTAGGA -3’), the PCR products were subcloned into a lentiviral backbone that expressing EGFP backbone driven by a CMV promoter, by seamless clone (NEB). For NDR2, PCR primers: FW: 5’- AGAATCCTGGCCCAGGATCCATGGCAATGACGGCAGG-3’, RV: 5’- ATAAGCTTGATATCGAATTCTCATAACTTCCCAGCTTTCATGTAG-3’. The pLV-Ubc-Pard3-GFP-WPRE was constructed in two steps: Pard3 was PCR-amplified and tagged with EGFP, then Pard3-EGFP was PCR-amplified using specific primers (5’- CACTTGGCGGCCGCCGCCACCATGGAACAGAAACTCATCTCTGAAG-3’ and RV: 5’- GAGGTTGATTAGGATCTATCGATTCATTACTTGTACAGCTCGTCCA-3’). The product was subcloned into a lentiviral vector under the Ubc promoter. The relative mutants were created by Site-directed mutagenesis, including NDR1 K118A, NDR2 K119A and Pard3 S144A. For the Tet-pLKO-puro inducible constructions (original from Addgene # 21915), the vector was a gift from Dr. David B. Lombard (University of Miami). The shRNA primer was synthesized and cloned into the backbone, which was digested with Age I and EcoR I. For the 6XHis-NDR1, 6XHis-NDR2and N-Pard3-6XHis used in kinase assay, PCR amplified 6XHis-NDR1 and 6XHis-NDR2 were subcloned into pcDNA 3.1 backbone. The N-Pard3-6XHis constructs were created by PCR amplifying N-Pard3 and subcloned into pET22B vector backbone between NdeI and SalI.

### Western blot

After experimental treatments, cells were washed three times with PBS and lysed in ice-cold radioimmunoprecipitation assay (RIPA) buffer (RIPA buffer, 150 mM sodium chloride, 1.0% NP-40, 0.5% sodium deoxycholate, 0.1% SDS, 50 mM Tris, pH 7.4 and 1mM EDTA) containing 1X Proteinase and Phosphatase inhibitor (Fisher Sci, PI78440). The protein concentration was measured by BCA Protein Assay (ThermoFisher, 23227). The protein samples (30μg) were prepared in Laemmli sample buffer (Bio-Rad, 1610747) and bolded for 5 minutes at 98℃ for denaturing. The prepared protein samples were separated on SDS-PAGE gel and transferred to the Nitrocellulose Membranes (Bio-Rad, 1620115). The membranes were blocked with blocking buffer (Rockland, RLMB-070-003) for 1 hour and incubated with primary antibody overnight at 4℃. The primary antibodies, including mouse anti-STK38 (NDR1) (1: 2000, Abnova, Cat# H00011329-M11), mouse anti-STK38L (NDR2) (1:2000, Origene, TA505176s), rabbit anti- Pard3(1:2000, Proteintech, 11085-1-AP). To control the loading, the membranes were immunoblotted with rat anti-alpha Tubulin (1:500, ThermoFisher, MA1- 80189) or mouse anti-beta Actin(1: 4000, Abcam, ab8224). After washing with TBST for 3 times, the membranes were incubated with secondary antibodies at room temperature for 2 hours with shaking, including IRDye® 680RD Goat-anti-Mouse IgG (1:15000, LI-COR, 926-68070), IRDye® 800CW Goat anti-Mouse IgG(1:15000, LI-COR, 926-32210), IRDye® 800CW Goat-anti-Rabbit IgG (1:15000, LI-COR, 926-32211), IRDye® 680RD Goat anti-Rat IgG(1:15000, LI-COR, 926-68076), Goat anti-Mouse IgG (H+L)-HRP (1:5000, ThermoFisher, 31430), Goat anti-Rabbit IgG antibody (1:4000, CST, 7074P2). The blots images were obtained using an Odyssey® CLx Imaging System (LI-COR) or Bio-Rad ChemiDoc Imaging System using ECL substrate (ACRES, 240329). Specific bands were quantified with Image Studio (LICORbio) or Image Lab software (BioRad) and the results were shown as the ratio of the intensity of loading control.

### Immunofluorescence

Cells were seeded onto coverslips pre-coated with 50 μg/mL Poly-D-lysine and subjected to the indicated treatment. After washing with PBS, cells were fixed with 4% paraformaldehyde (PFA) for 10 minutes and blocked in PBS containing 0.1% Triton X100, 1% of BSA and 0.1 M glycine for 2 hours at room temperature. After blocking, the slides were incubated with primary antibodies overnight 4℃ with antibodies including mouse anti-Vinculin (1: 300, Sigma, V9131), rabbit anti-GM130 (1:1000, CST, 12480T), rabbit anti- Pard3(1:400, Proteintech, 11085-1-AP), mouse anti-Zo-1 (1:500, Proteintech, 66452-1-Ig). Next day, the slides were washed with PBS 3 times/5minutes with gentle shaking. Then, the slides were incubated with secondary antibodies that were conjugated with Alexa Fluor dye (488, 555 or 647) for 2 hours at room temperature. After incubation with secondary antibodies, the slides were applied to DAPI (0.5 μg/mL) staining for 10 minutes. The slides were imaged with a Zeiss Axio Observer inverted microscope.

### Live cell Cdc42 biosensor imaging

The HFF cells were stably expressed Cdc42 localization-based biosensor (dimeric Tomato-WASp (CRIB), Addgene #176099) via lentivirus infection(Mahlandt et al., 2023). The cells express NDR kinases shRNA (Scramble as control) were further stable expressed with Cdc42 biosensor (dTomato as fluorescent marker), and FACS was further utilized to enrich dual positive cells. After several generation amplification, the cells were applied for visualization of active Cdc42 via a Zeiss Axio Observer inverted microscope with an incubation equipment for automated live-cell imaging. The cells were maintained in 50% of Dulbecco’s modified Eagle’s medium and 50% of Ham’s F- 10 Nutrient Mix (Gibco, 10569-010, 11550-043) with 10% FBS. For imaging experiment, the cells were plated on Cellview cell culture dish with glass bottom (Greiner) the day before imaging. Imaging was performed in culture medium (50% DMEM+50% Ham’s F- 10) supplemented with 10mM HEPES and 10% of FBS in a heated chamber with a lid (maintained at 37℃). The time-lapse sequences were obtained by a Zeiss Axio Observer inverted microscope equipped with an objective Plan-Apochromat 63x/1.4 Oil M27 lens, PvCam camera, Zeiss Zen 3.4 Pro software. The images were recorded with an exposure time for dTomato with 1 s and taken at 10s intervals for a total 1000 s recording time(Machacek et al., 2009).

### Cell edge tracking and windowing for imaging analysis

For cell-edge tracking and windowing analysis, we used a custom MATLAB toolbox, u- register, originally developed in the Danuser Lab (Machacek et al., 2009). This software allows the analysis of local cell edge motions, including protrusion and retraction, through automated windowing and signal analysis at the cell boundary. The conception of ‘windows’ refers to dividing the cell interior into sampling regions, each associated with a nearby segment of the cell edge based on proximity. Briefly, the cell edges were segmented by applying intensity thresholding to the dTomato signal, and the edge displacements were tracked using a mechanical model previously described in the literature (Machacek and Danuser, 2006). Multi-Scale-Automatic (MSA) segmentation was applied to analyze edge motion profiles (Noh et al., 2022), enabling the measurement of edge velocities and Cdc42 activity from segmented cell boundaries in live-cell imaging. We used the ‘Constant number’ option to propagate the windows, maintaining a constant perpendicular width of 1.02 μm at each time point. As the length of the cell boundary changed over time, the size of individual windows varied accordingly, while the total number of windows remained constant. This window-based segmentation method allowed the extraction of time series data for local edge velocity and Cdc42 activity signals for further analysis.

### Cross-correlation analysis of edge velocity and Cdc42 biosensor intensity

To quantify the temporal relationship between edge velocity and Cdc42 GTPase activity, we performed a pearson’s correlation coefficient analysis between two activity time series using MATLAB *xcov* function (Machacek et al., 2009). For *x*(*t*) be the Cdc42 activity and *y*(*t*) be the edge velocity measured at discrete time points t = 1, 2 , … ,T.

For each time series pair (x(t), y(t)), [c, lags] = xcov(x, y, maxlag, ’normalized’) was computed. We employed the ‘normalized’ option to obtain a correlation coefficient ranging between -1 and +1. For two signals *x* and *y*, the **normalized cross-correlation** at lag *k* is defined as:

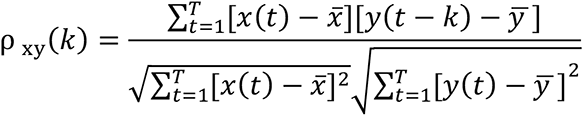

Where *x̅* and ȳ are the means value of *x* and *y*, respectively, and *k* is the time lag. We extracted two metrics: 1. Peak correlation coefficient, and 2. Time lag at peak for further analysis. The correlation coefficient of all sampling windows of a cell was averaged by a spline fit (by Origin software) for a better estimate of the average timing between edge velocity and Cdc42 GTPase activity. The samples were eliminated from further analysis if the correlation maximum/minimum did not reach above/below the 95% confidence levels of the Pearsons’s correlation coefficient. The 95% confidence level of Pearson’s correlation coefficient is calculated in function *CONFIDENCE* in excel software.

### Scratch wound healing and Migration behavior assays

Cell wound healing and migration assays were performed on confluent cell layers for different treatments. The cell migration was initiated by a scratch on monolayers vertically with a 200 µL pipette tip. Thirty minutes after scratching, cells were washed twice with warmed DMEM and addition of complete medium. For Doxycycline(Dox) induced NDR kinases knock-down, the cells were pre-treated with/without 5ng/mL or 10ng/mL of Dox 3 days before splitting. Then the cells were cultured in the medium contains Dox to be confluent for scratching assay in a 96-well plate for behavior testing or 24-well plate on PDL treated coverslips for immunofluorescent staining 10 h after scratching. Live cell migration behaviors were recorded using an Incucyte ® SX5 Live-Cell Analysis System(Sartorius). The Incucyte platform allows for long-term, time-lapse experiments under physiological conditions, which can maintain temperature, humidity, and CO_2_ levels to support normal cell function. High-resolution images were automatically acquired at defined intervals (e.g., every 15 or 20 min) over the course of the experiment, enabling real-time kinetic monitoring of cell migration. The Incucyte software (2023A Rev2) simultaneously analyzed the captured images, allowing for quantitative confluence in real time. This automated and continuous monitoring minimized experimental variability and provided immediate insights into dynamic changes in cell migrating behavior.

### Rho GTPase activity assay

The plasmids (pGEX-2T backbone) containing the coding sequences of PAK (GST-PBD, amino 23-137) for Cdc42/Rac-interaction binding (CRIB) motif and Rhotekin (GST-RBD, amino acids 7-89, addgene # 15247) for Rho-binding domain were used for affinity-precipitation of cellular GTP bounded Cdc42/Rac and RhoA, respectively. The recombinant proteins were bacterially expressed in BL21 E. Coli (a gift from Dr. Daniel G. Isom, University of Miami) with an auto-induction system (Grabski et al., 2005).

Bacteria cells were collected by centrifugation and washed with PBS twice, followed by sonication in PBS supplied with 1X Proteinase and Phosphatase inhibitor (Fisher Sci, PI78440). After centrifuge, MagneGST^TM^ Glutathione Particles (Promega, V8611) were used to clarified lysates and incubated 3 hours at 4 ℃ with rotation. GST-PBD and GST-RBD beads were washed with PBS with proteinase inhibitor and used in Rho GTPase activity assay immediately.

HFF cells with 70-80% confluency were washed with PBS and lysed in cold non-denatured immunoprecipitation buffer (described above) supplied with 10 mM MgCl2 and 1X Proteinase and Phosphatase inhibitor (Fisher Sci, PI78440) and 0.2 mM PMSF in ice for 30min. Total protein concentration was measured by BCA Protein Assay (ThermoFisher, 23227). About 800-1000 μg of total protein was incubated with 50μl of GST-PBD or GST-RBD beads at 4 ℃ overnight with rotation. The beads were washed with lysis buffer for 5 times and eluted with 2X Laemmli Sample Buffer at 98℃ with shaking for 10 minutes. The Cdc42, Rac and RhoA proteins were detected by western blotting using a mouse anti-Cdc42 (1:200, Santa Cruz, sc-8401), mouse anti-RhoA (1:500, Santa Cruz, sc-418), mouse anti- Rac 1/2/3( 1:500, Santa Cruz, sc-514538).

The results were analyzed as described as western blot pipeline.

### Kinase assay

Recombinant 6XHis-NDR1 and 6XHis-NDR2 protein were expressed in HEK293T cells by transient transfection for 48 hours, then the kinases were lysed in IP-buffer (20mM Tris-HCl pH 8.0, 150mM NaCl, 1% NP-40, 50mM NaF, 1 mM phenylmethylsulfonyl fluoride (PMSF), 1X protease and phosphatase inhibitor cocktail (Halt Protease and Phosphatase Inhibitor Cocktail (100X))) and purified by Ni-NTA Magnetic Agarose Beads (QIAGEN) according to the protocol. Briefly, one 10-cm dish cells were lysed in 600μL non-denature IP-buffer followed by 40 μL Ni-NTA Magnetic Agarose Beads, rotate at 4℃ for 2 hours, washed twice with 500 μL lysate buffer (150mM NaCl), one time with 500 μL lysate buffer (1M NaCl), one time with 500 μL lysate buffer (150mM NaCl), and then washed one time with 500 μL kinase buffer (50mM Tris-HCl, pH 7.5, 100mM NaCl, 10mM MgCl2, 1mM MnCl2). The beads were resuspended in 125 μL kinase buffer for further use. 6XHis-tagged recombinant N-terminal truncated N-Pard3- WT and N-Pard3-S144A mutant were bacterial expressed by IPTG induction (DE3 cells) and were lysed in 1X PBS, followed by further Ni-NTA Magnetic Agarose Beads purification. 10 μL of beads with bacterially expressed N-Pard3-6X his protein, 25 μL of beads with NDR1 or NDR2 protein, 4 μL of 1mM ATPγS were mixed and incubated for 60min at 30 ℃ and stopped with adding 20mM EDTA, and alkylated by incubating with 2.5 mM p-nitrobenzylmesylate (Sigma-Aldrich, stock 50 mM) for 2 h. Then, 17 μL of 4X Laemmli buffer was added and boiled at 98 ℃ for 3min. The sample was analyzed by western blot, and thiophosphorylation of N-Pard3 was detected by thiophosphate ester antibody (Epitomics), total N-Pard3 with Myc antibody (Santa Cruz Biotechnology) and His-antibody (Santa Cruz Biotechnology), NDR1 and NDR2 were stained with anti- NDR1 and anti-NDR2 antibody, respectively.

### Human skin ex vivo wound healing model

Health human skin samples were applied to generate acute wounds as described in previous publications (Jozic et al., 2019; Marjanovic et al., 2022; Sawaya et al., 2019). The acute wounds were generated under sterile conditions. The subcutaneous fat was removed prior to a 3-mm punch for making wound in the epidermis. Then the skin discs (8-mm) with 3-mm epidermal wound in the center were collected by an 8-mm biopsy punch, and the samples were infected with lentivirus carrying shRNA (∼ 1X 10^7^ Tu/ml, 100μl + 10μg/ml Polybrene) in 1ml of fresh media for 6 hours. After 6-hour infection, the samples were washed with PBS for 5 times and transferred to the air-liquid interface and maintained at the air-liquid interface at 37℃ with 5% CO2 incubator for 4 days. The air-liquid interface was changed with fresh media every 24 hours. Wounds were collected on day 4 after wounding. The protein levels were confirmed by western blot.

The wound healing was evaluated by H&E staining.

### H&E staining and immunofluorescent staining from Paraffin sections

Wounded skin specimens were fixed in 10 % neutral-buffered formalin overnight at room temperature, then processed for paraffin embedding and microtome sectioning. The sliced specimens were under deparaffinization and rehydration. For H&E staining, tissue sections were stained with hematoxylin and eosin following established protocols (Feldman and Wolfe, 2014). For immunofluorescent staining, the tissue sections were under antigen retrieval by heating in Tris EDTA pH 9.0 + 0.5% Tween20 at 95 ℃ for 30min followed with 20min cooling in ice-cold water. Then, the tissue specimens were rinsed 3 times in TBS and blocked with background punisher (BioCare Medical, REF BP9744) and incubated with primary antibody, mouse anti-cytokeratin 17 (1:250, Santa Cruz, SC-393091) overnight at 4℃. Next day, the slides were rinsed 3 times/5min in 1X TBST, followed by secondary antibody incubation for 1 hour at room temperature. The fluorescence slides were imaged with a Zeiss 880 confocal microscope.

### Microscopy and image analysis

Fluorescence images of cells or tissues were taken on either a Zeiss Axio Observer inverted microscope equipped with PvCam Camera(Zeiss) or Zeiss 880 confocal microscope (Zeiss) at 22℃ with 10X (NA:0.3), 40X (NA:0.6) and 63X (NA:1.4) Plan-Apochromat objectives. All the data was acquired using ZEN imaging suite (Zeiss). The image analysis, including morphometrics, intensity processing, edge velocity measurement, and individual cell migrating tracking, were performed using ZEN imaging software, opera phenix plus High-content screening system, MATLAB software, RStudio, Python, Incucyte software (2023A Rev2) and an open access National Institutes of Health (NIH) software platform, FIJI.

The live cell migrating behavior tracking was performed using a semiautomated tracking plugin, TrackMate in FIJI software. The raw data obtained from FIJI software was analyzed using program R code in RStudio software and visualized by MATLAB, OriginPro 2021 and GraphPad Software.

## Statistics

Sample sizes and experimental designs were determined by power analysis or informed by comparable studies in literature. All the statistical tests were two-tailed. When only two groups were compared, an unpaired Student’s T-test was applied. For three or more groups, one-way ANOVA was used; if the difference was considered significant, Bonferroni’s post-hoc test identified pairwise differences. The cell migration/wound healing data were analyzed with two-way repeated-measures ANOVA, followed by Tukey’s post-hoc comparisons between time points if differences were significant. All analyses were performed in GraphPad Prism 9.5.1 (GraphPad Software, Inc.). Data was presented as mean ± SEM, with statistical significance set at *p*-value < 0.05 ( * P < 0.05, ** P < 0.01, *** P < 0.001).

## Acknowledgments

We would like to thank Kannappan Sriramajayam and Ganapathi Kandasamy for their technical assistance, Zhipeng Meng for insightful discussions and technical support, Daniel Isom for providing E.coli DE3 cells, Kerry Burnstein for sharing Incucyte recording system, Qiulei Li for assistance in developing the program code used in this study, and Laura Doyle for assistance in manuscript writing.

Research reported in this publication was supported by the National Institutes of Health R01 Grant no. R01GM129514 and by the University of Miami Sylvester Comprehensive Cancer Center which receives funding from the National Cancer Institute (NCI) of the National Institutes of Health (NIH) under award number P30CA240139. The content is solely the responsibility of the authors and does not necessarily represent the official views of the NIH. The salary of FZ is in part supported by R01GM138557.

The authors declare no competing financial interests.

## Author contributions

Conceptualization: J. Gu., F. Zhang., and F. Verde.; Investigation: J. Gu., J. Marjanovic.; Funding acquisition: F. Verde., F. Zhang., and M. Tomic-Canic.; Supervision: F. Verde.; Writing – Original draft: J. Gu.; Writing- review & editing: J. Gu., J. Marjanovic., M. Tomic-Canic., F. Zhang., and F. Verde.

