## Supplementary Figures 1-6 for "NDR1/2 kinases regulate cell polarization and cell motility through Cdc42 GTPase and Pard3 signaling in mammalian cells"

**NDR1/2 kinases regulate persistent cell motility through Pard3-Cdc42 signaling in migrating fibroblasts**

**The file includes:**

Figure S1. **NDR1 and/or NDR2 depletion delays 2-D cell wound closure speed.**
Figure S2. **Cells display no Golgi polarization in random monolayers.**
Figure S3. **NDR kinase knock down changes Rac and RhoA GTPases activity.**
Figure S4. **Pard3 expression and location in Tet-inducible NDR1 or NDR2 knockdown system.**
Figure S5. **The validation of Pard3 and NDR1/2 plasmid constructions.**
Figure S6. **NDR1 protein expression by western blot in Tet-on inducible NDR1 knockdown in HaCaT cells**

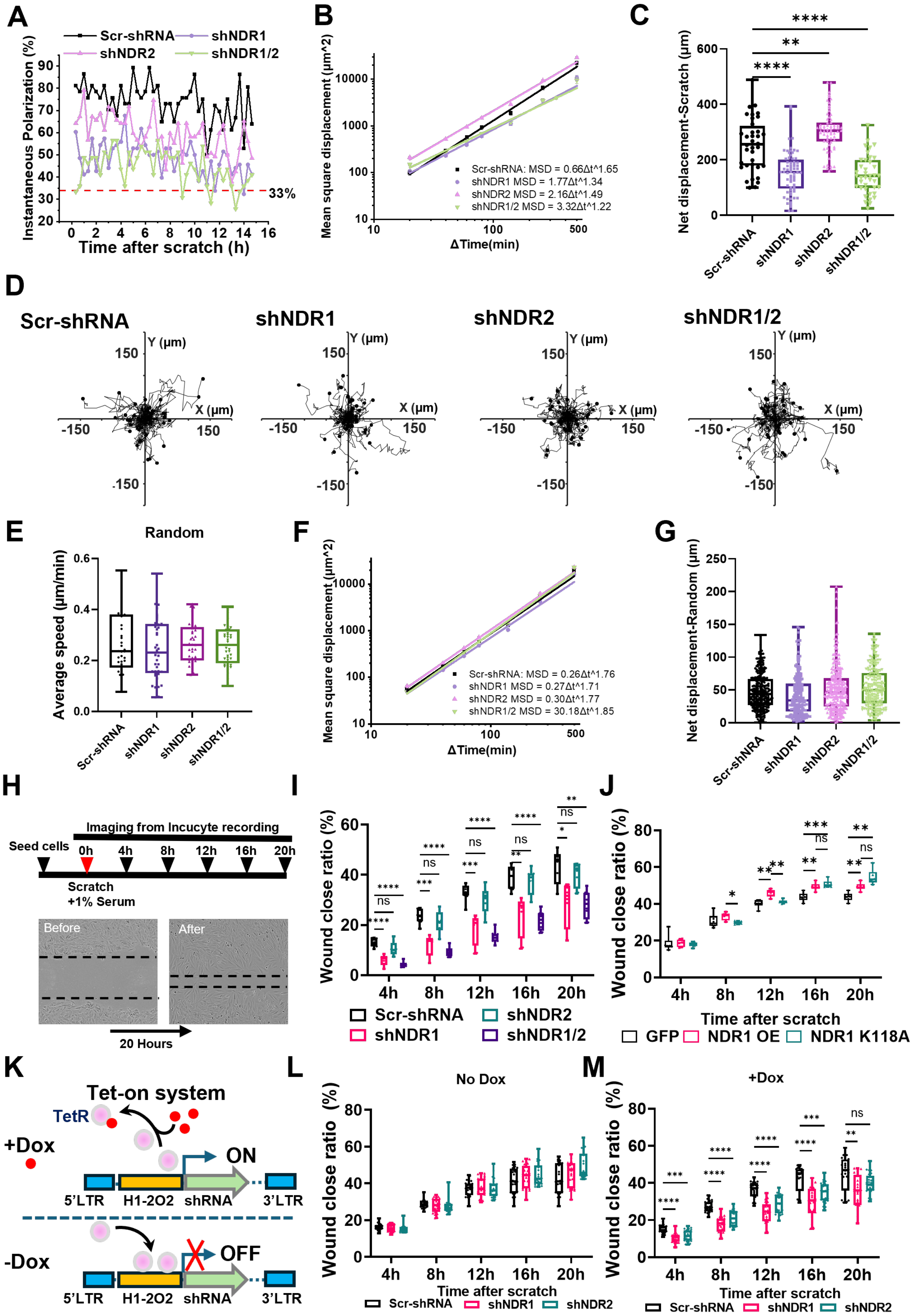
A
Scr-shRNA
shNDR1
shNDR2
shNDR1/2
Instantaneous Polarization (%)
33%
Time after scratch (h)
B
Mean square displacement (μm^2)
Scr-shRNA: MSD = 0.66Δt^1.65
shNDR1 MSD = 1.77Δt^1.34
shNDR2 MSD = 2.16Δt^1.49
shNDR1/2 MSD = 3.32Δt^1.22
ΔTime(min)
C
Net displacement-Scratch (μm)
D
Scr-shRNA
shNDR1
shNDR2
shNDR1/2
Y (μm)
X (μm)
E
Random
Average speed (μm/min)
F
Mean square displacement (μm^2)
Scr-shRNA: MSD = 0.26Δt^1.76
shNDR1 MSD = 0.27Δt^1.71
shNDR2 MSD = 0.30Δt^1.77
shNDR1/2 MSD = 30.18Δt^1.85
ΔTime(min)
G
Net displacement-Random (μm)
H
Imaging from Incucyte recording
Seed cells
0h 4h 8h 12h 16h 20h
Scratch
+1% Serum
Before
After
20 Hours
I
Wound close ratio (%)
Scr-shRNA
shNDR1
shNDR2
shNDR1/2
J
Wound close ratio (%)
Time after scratch
GFP
NDR1 OE
NDR1 K118A
K
Tet-on system
TetR
+Dox
ON
5'LTR H1-2O2 shRNA 3'LTR
-Dox
OFF
L
No Dox
Wound close ratio (%)
Time after scratch
Scr-shRNA shNDR1 shNDR2
M
+Dox
Wound close ratio (%)
Time after scratch
Scr-shRNA shNDR1 shNDR2

Figure S1. **NDR1 and/or NDR2 depletion delays 2-D cell wound closure speed. (A)** Instantaneous cell polarization in leading cells. The data points > 30 $^{o}$ were defined as polarity cells. 33% is the random cell polarity that indicates the cell orienting itself in one of the three possible outcomes by chance. **(B)** The mean square displacement of cells after scratch, Scr-shRNA ($R^2$ = 0.98), shNDR1 ($R^2$ = 0.96), shNDR2 ($R^2$ = 0.98), shNDR1/2 ($R^2$ = 0.96). **(C)** Net displacement of scratch monolayers over 15h. ****P <0.0001, **P<0.01; One-Way ANOVA followed by Bonferroni's test, n = 37-69. **(D)** Representative trajectories of fibroblasts seeded at low densities for 15h. **(E)** Average random movement speed per cell over 15 h. **(F)** The mean square displacement of cells with random movement, Scr-shRNA ($R^2$ = 0.98), shNDR1 ($R^2$ = 0.97), shNDR2 ($R^2$ = 0.98), shNDR1/2 ($R^2$ = 0.98). **(G)** Net displacement of random cells over 15h. **(H)** Schematic of time series cell wound closure measurement by Incucyte Imaging recording system. **(I-J)** Wound closure ratio. ***P <0.001, **P<0.01, *P<0.05; Two-Way ANOVA followed by Tukey's post hoc tests. n = 8. **(K)** Schematic of pLKO-Tet-On system. **(L-M)** Wound closure ratio with an inducible knock down system. HFF cells were stabilized expressed with pLKO-Tet-on lentivirus shRNA targeting *Ndr1* or *Ndr2*. A wound closure ratio measurement was performed, and the time series images were captured by Incucyte real-time imaging system. ****P <0.0001, ***P <0.001, **P<0.01; Two-Way ANOVA followed by Tukey's post hoc tests. n = 24.

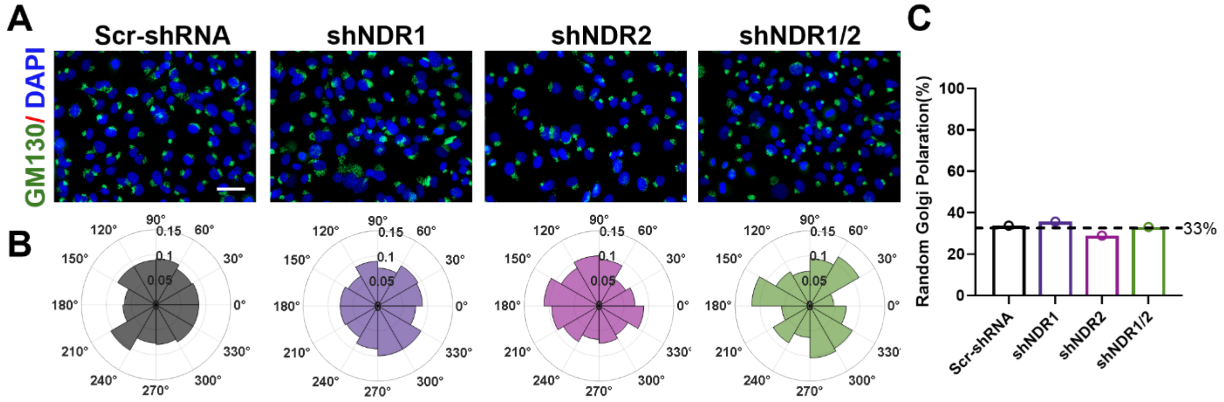


Figure S2. **Cells display no Golgi polarization in random monolayers. (A)** Golgi apparatus (GM130-Alexa Fluor 488) and nuclear (DAPI) were visualized, Bar = 50 μm. **(B)** Rose plot of Golgi polarization in Random cells in Scr-shRNA (Dark), shNDR1(Medium purple), shNDR2 (Vibrant purple) and shNDR1/2 (Green). (C) Quantification of Golgi orientation in random cells.

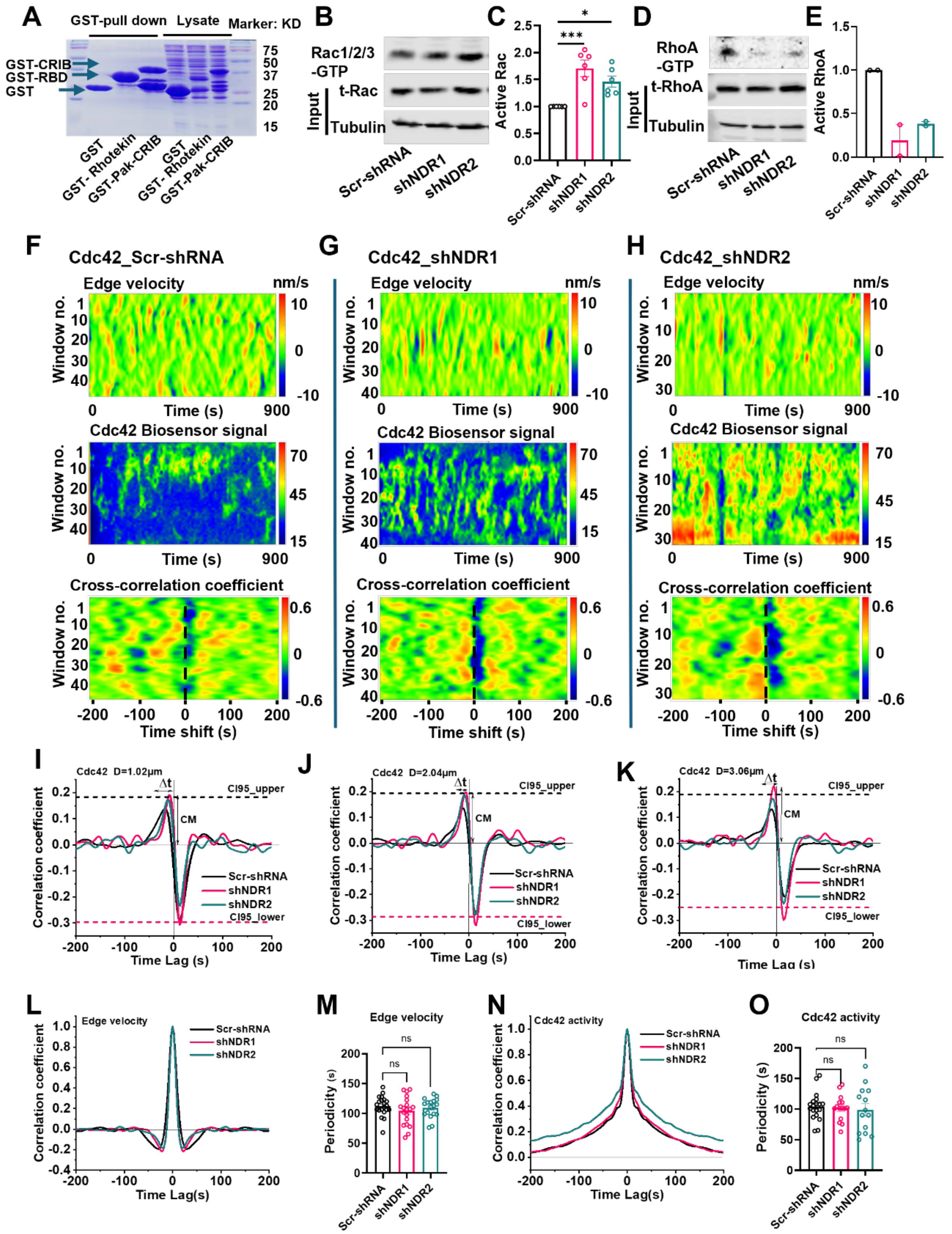


Figure S3. **NDR kinase knock down changes Rac and RhoA GTPases activity. (A)** Coomassie Blue staining of Rhotekin and Pak-CRIB proteins expressed in E. coli DE3

cells. **(B-E)** Western blot analysis of active Rac (B–C) and active RhoA (D–E) detected by GST pull-down assays using GST–Pak-CRIB or GST–RBD, respectively, from lysates of NDR1- or NDR2-knockdown HFF cells; ***$P<0.001$, *$P<0.05$, One-Way ANOVA followed by Bonferroni's test with Scr-shRNA, n= 5; **(F-H)** Representative time-course maps of cellular edge dynamics, Cdc42 activity, and cross-correlation coefficients between edge velocity and Cdc42 GTPase signaling at defined subcellular distances (D = 1.02–2.04 μm). **(I-K)** Cross-correlation coefficients between edge velocity and Cdc42 GTPase activity measured at distances of D = 1.02, 2.04, and 3.06 μm from the cell edge.

**(L-O)** Temporal autocorrelation of edge velocity (L) and Cdc42 (N) fluctuations measured at the leading edge. The periodicity of edge velocity (M) and Cdc42 activity (O) were quantified, with no significant differences observed (ns; n = 18–21).

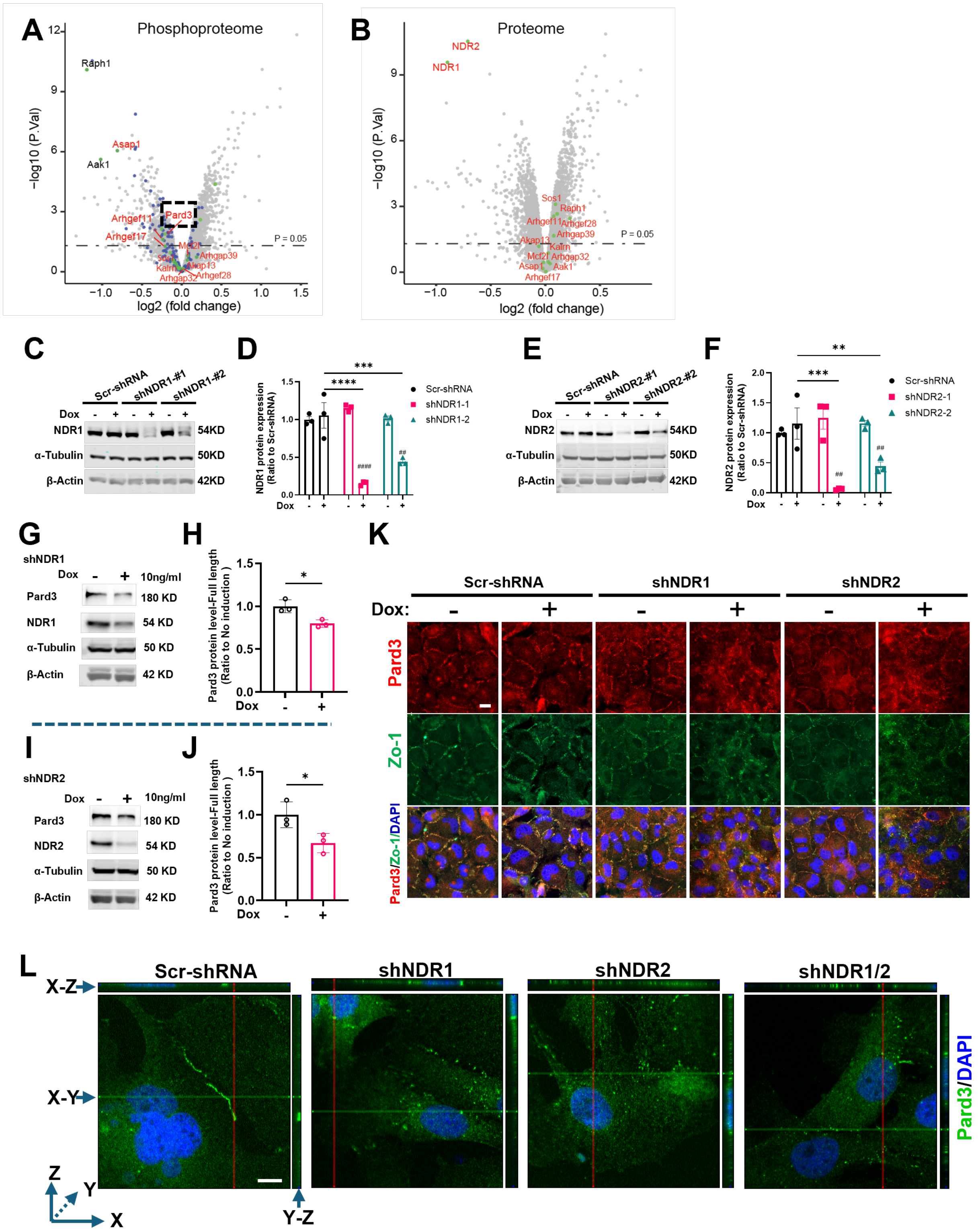

Figure S4. **Pard3 expression and location in Tet-inducible NDR1 or NDR2 knockdown system. (A-B)** Volcano plot of phosphopeptide and protein levels between control and NDR1/2 knockout group in mice hippocampus. The blue dots represent phophopeptides with NDR consensus motif (H.R..[ST]). **(C-F)** Western blot confirms the Tet-on inducible shRNA knock-down effect of NDR1(D-E) or NDR2 (F-G) protein level, ****P <0.0001, ***P <0.001, **P<0.01; Two-Way ANOVA followed by Tukey's post hoc tests; ####P<0.001, ##P<0.01, student's T-test, n = 3. **(G-J)** Doxycycline-induced knockdown of NDR1 or NDR2 resulted in reduced Pard3 protein expression, as confirmed by western blot analysis. *P <0.05, two-tailed unpaired Student's t test, n=3. **(K)** Cells were treated with or without doxycycline for 3 days, fixed with 4% PFA, and subjected to immunofluorescence analysis. Pard3 and ZO-1 co-localization in fibroblasts were examined using antibodies against Pard3 and ZO-1, visualized with Alexa Fluor 647 and Alexa Fluor 488, respectively. Nuclei were counterstained with DAPI. Bar = 20 μm. (L) Confocal image 3-D reconstruction for the analysis of Pard3 cellular location, Pard3 was examined using antibodies against Pard3 visualized with Alexa Fluor 488. Bar = 10 μm.

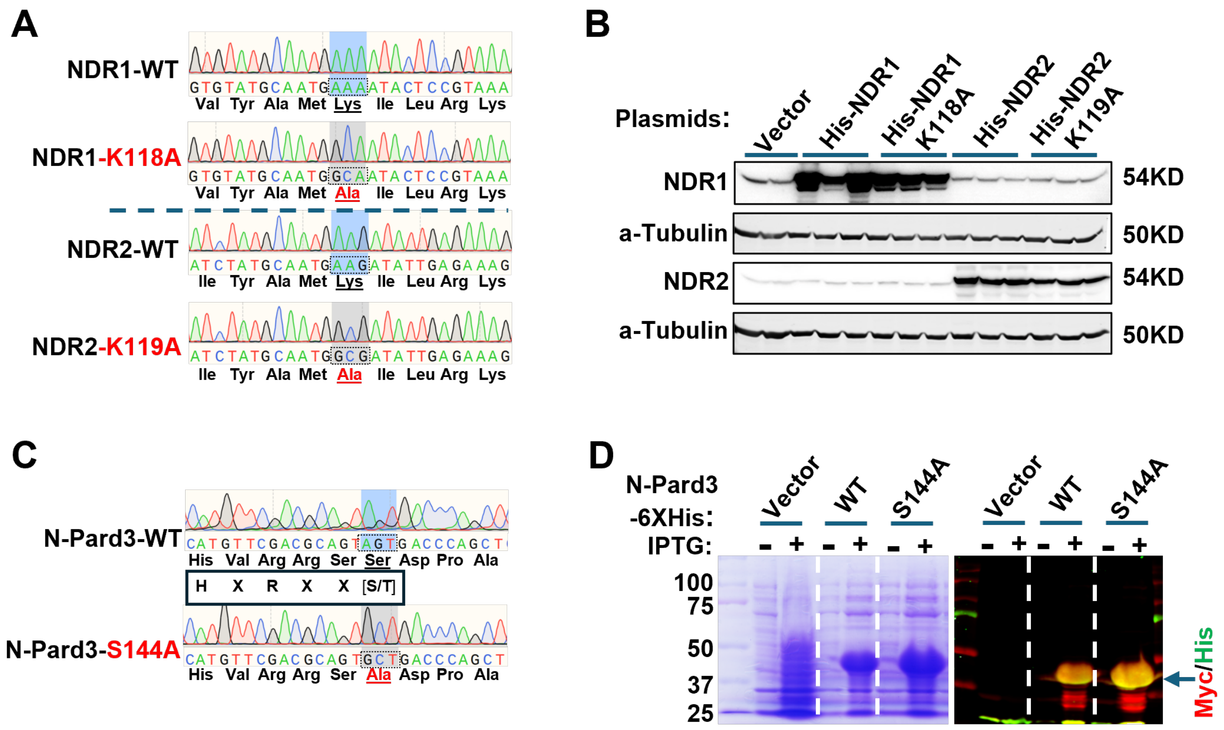


Figure S5. **The validation of Pard3 and NDR1/2 plasmid constructions. (A,C)** Plasmid constructs of NDR1, NDR2, and Pard3, including their respective mutants, were validated by Sanger sequencing. **(B)** Western blot confirms the expression of His-NDR1 and His-NDR2 with their kinase dead mutation expression in HEK293T cells. **(D)** Coomassie Blue staining (left) of N-Pard3 peptides which were induced by IPTG (0.5mM) in E.coli DE3 cells. Western blot shows the N-Pard3 peptides expression by probe with both Myc(Red) and His(Green) antibody.

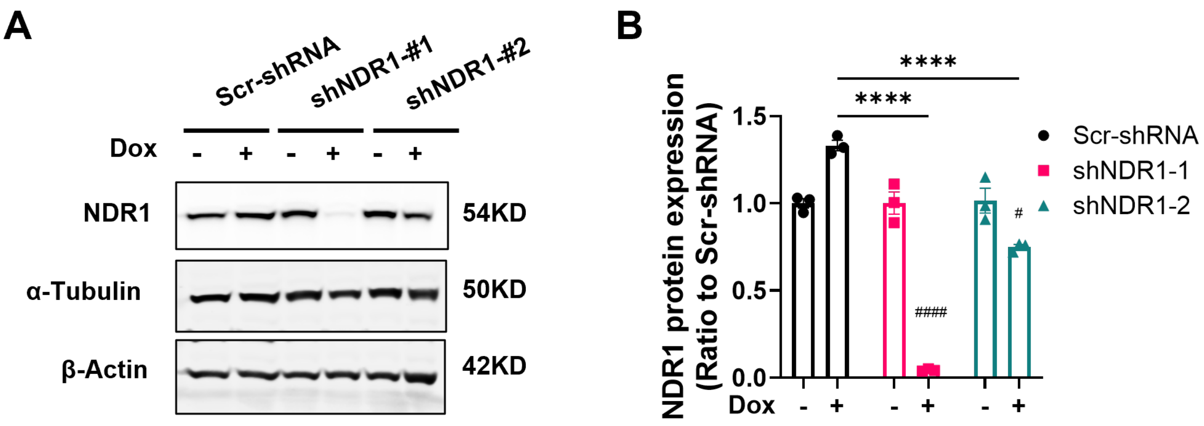


Figure S6. **NDR1 protein expression by western blot in Tet-on inducible NDR1 knockdown in HaCaT cells. (A-B)** Western blot confirms the expression of NDR1 protein expression in Dox-induced knock down HaCaT cells.
